# Removal of high frequency contamination from motion estimates in single-band fMRI saves data without biasing functional connectivity

**DOI:** 10.1101/837161

**Authors:** Caterina Gratton, Ally Dworetsky, Rebecca S. Coalson, Babatunde Adeyemo, Timothy O. Laumann, Gagan S. Wig, Tania S. Kong, Gabriele Gratton, Monica Fabiani, Deanna M. Barch, Daniel Tranel, Oscar Miranda-Dominguez, Damien A. Fair, Nico U. F. Dosenbach, Abraham Z. Snyder, Joel S. Perlmutter, Steven E. Petersen, Meghan C. Campbell

## Abstract

Denoising fMRI data requires assessment of frame-to-frame head motion and removal of the biases motion introduces. This is usually done through analysis of the parameters calculated during retrospective head motion correction (i.e., ‘motion’ parameters). However, it is increasingly recognized that respiration introduces factitious head motion via perturbations of the main (B0) field. This effect appears as higher-frequency fluctuations in the motion parameters (> 0.1 Hz, here referred to as ‘HF-motion’), primarily in the phase-encoding direction. This periodicity can sometimes be obscured in standard single-band fMRI (TR 2.0 – 2.5 s.) due to aliasing. Here we examined (1) how prevalent HF-motion effects are in seven single-band datasets with TR from 2.0 - 2.5 s and (2) how HF-motion affects functional connectivity. We demonstrate that HF-motion is relatively trait-like and more common in older adults, those with higher body mass index, and those with lower cardiorespiratory fitness. We propose a low-pass filtering approach to remove the contamination of high frequency effects from motion summary measures, such as framewise displacement (FD). We demonstrate that in most datasets this filtering approach saves a substantial amount of data from FD-based frame censoring, while at the same time reducing motion biases in functional connectivity measures. These findings suggest that filtering motion parameters is an effective way to improve the fidelity of head motion estimates, even in single band datasets. Particularly large data savings may accrue in datasets acquired in older and less fit participants.

**Highlights:**

- Single-band fMRI motion traces show factitious high-frequency content (*HF-motion*)
- The magnitude of HF-motion relates to age and other demographic factors
- HF-motion elevates framewise displacement (FD) and causes data loss
- Substantial fMRI data can be recovered from censoring by filtering motion traces
- Filtering motion traces reduces motion artifacts in functional connectivity

## Introduction

Functional connectivity (FC) measured with functional MRI (fMRI) offers a powerful means of examining the systems-level organization of the human brain (Biswal et al., 1995; Buckner et al., 2013; Power et al., 2014b). However, the last decade has seen an explosion of studies demonstrating that numerous sources of artifacts systematically distort estimates of FC, including head motion (Power et al., 2012; Satterthwaite et al., 2019; Satterthwaite et al., 2012; Van Dijk et al., 2012) and respiration (Birn, 2012; Birn et al., 2008; Power et al., 2017b). Head motion introduces bias in measured FC through both common effects across all pair-wise regional correlations as well as distance-dependent biases, where correlations are increased most for adjacent regions and relatively decreased for regions that are distant (Power et al., 2012; Satterthwaite et al., 2012). These distortions are of particular concern in studies comparing groups or conditions that differ systematically in head motion, for example in comparisons between children and young adults (Greene et al., 2016; Nielsen et al., 2019; Satterthwaite et al., 2013), adults of different ages (Madan, 2018; Savalia et al., 2017), or clinical populations vs. normative controls (Dosenbach et al., 2017; Fair et al., 2012; Gratton et al., 2019). Fortunately, effective approaches have been developed to reduce biases introduced by head motion (see review by (Power et al., 2015); (Ciric et al., 2017; Parkes et al., 2018)). One common approach is to use a combination of global signal regression (GSR) and motion censoring (removal of BOLD volumes with high levels of motion) (Power et al., 2014a); these two steps in combination perform well in controlling the link between head motion and FC (Ciric et al., 2017; Parkes et al., 2018). Motion censoring is particularly efficacious at reducing distance-dependent artifacts in fMRI (Ciric et al., 2017; Parkes et al., 2018; Power et al., 2014a), but comes at the cost of reducing the amount of useable data (Ciric et al., 2017; Parkes et al., 2018; Raut et al., 2019).

Head motion in fMRI is typically estimated from a summary statistic of frame-to-frame motion (e.g., framewise displacement, FD (Power et al., 2012), or relative root-mean-square movement, RMS (Satterthwaite et al., 2013)) derived from retrospective functional image alignment (i.e., “motion correction”). However, realignment estimates may not be pure measures of head-motion and may be influenced by other factors such as respiration. Respiration may influence realignment estimates in two ways: (a) through true changes in head position caused by respiration-related movement and (b) apparent (factitious) motion in the phase-encoding direction generated by perturbations of the main (B0) magnetic field of T2* images caused by chest wall motion (Brosch et al., 2002; Chen et al., 2019; Durand et al., 2001; Fair et al., 2020; Power et al., 2019; Raj et al., 2001; Van de Moortele et al., 2002). Evidence suggests that factitious motion may be the predominant source of respiration-related motion artifacts in fMRI ((Brosch et al., 2002; Raj et al., 2001); elaborated on further in the discussion).

The rate of respiration in humans depends on age and cardiopulmonary status, but typically is 12-18 breaths per minute (0.2-0.3 Hz.) in adults (Charlton et al., 2018). This frequency range is above the Nyquist folding frequency of most single-band fMRI studies. Specifically, at an image sampling frequency of 0.4-0.5 Hz (i.e., TR from 2.0-2.5 s.), the Nyquist frequency is 0.2 – 0.25 Hz^1^, such that adult respiration rates would alias into frequencies from 0.1-0.2 Hz. Thus, respiratory effects on alignment estimates (factitious or real) are more clearly identifiable in head motion traces from typical multi-band acquisition sequences that allow for data collection at fast rates (TRs<1.5 s. have Nyquist limits > 0.33 Hz.) (Fair et al., 2020). This faster sampling allows investigators to separate respiratory-related effects from other head motion effects, as abrupt head motion effects are broadband, while gradual head motion shifts tend to occur at low frequencies. Indeed, recent evidence has demonstrated that fast multiband fMRI data exhibit systematic perturbations that match respiration rates (Fair et al., 2020) and are most prominent in the phase-encoding direction, consistent with an influence on magnetic field heterogeneity (Fair et al., 2020).

However, preliminary evidence from (Fair et al., 2020) suggests that respiration may also contaminate motion estimates from fMRI data collected at typical slower acquisition rates, in this case at aliased frequencies from 0.1-0.2 Hz. Here, we build on (Fair et al., 2020) to ask how strong this contamination is across datasets and participants with diverse characteristics. Importantly, it is also still uncertain how respiration-related content in motion parameters might affect functional connectivity, in light of common denoising techniques that censor even small movements. Thus, the two major goals of this investigation were (1) to determine the prevalence of motion at higher, respiration-related, frequencies (*HF-motion*, >0.1 Hz.) in single-band fMRI and (2) to determine whether HF-motion can be removed without adversely affecting functional connectivity. To this end, we analyzed data from seven single-band datasets collected on different populations on diverse scanners and with different sequences (**Table 1**). We examined participant characteristics associated with the occurrence of high-frequency motion, demonstrating that in slower TR datasets HF-motion is more prominent in older individuals and those with a higher body mass index (BMI). Finally, we propose a low-pass filtering solution to correct HF-motion contamination. We demonstrate that this solution saves substantial amounts of data from censoring while still effectively reducing motion-related bias in FC, following Ciric et al. (2017). These results have the potential to substantially increase data savings and increase the fidelity of head motion estimates, particularly in datasets focused on older adults or those with higher BMI.

**Table 1:**
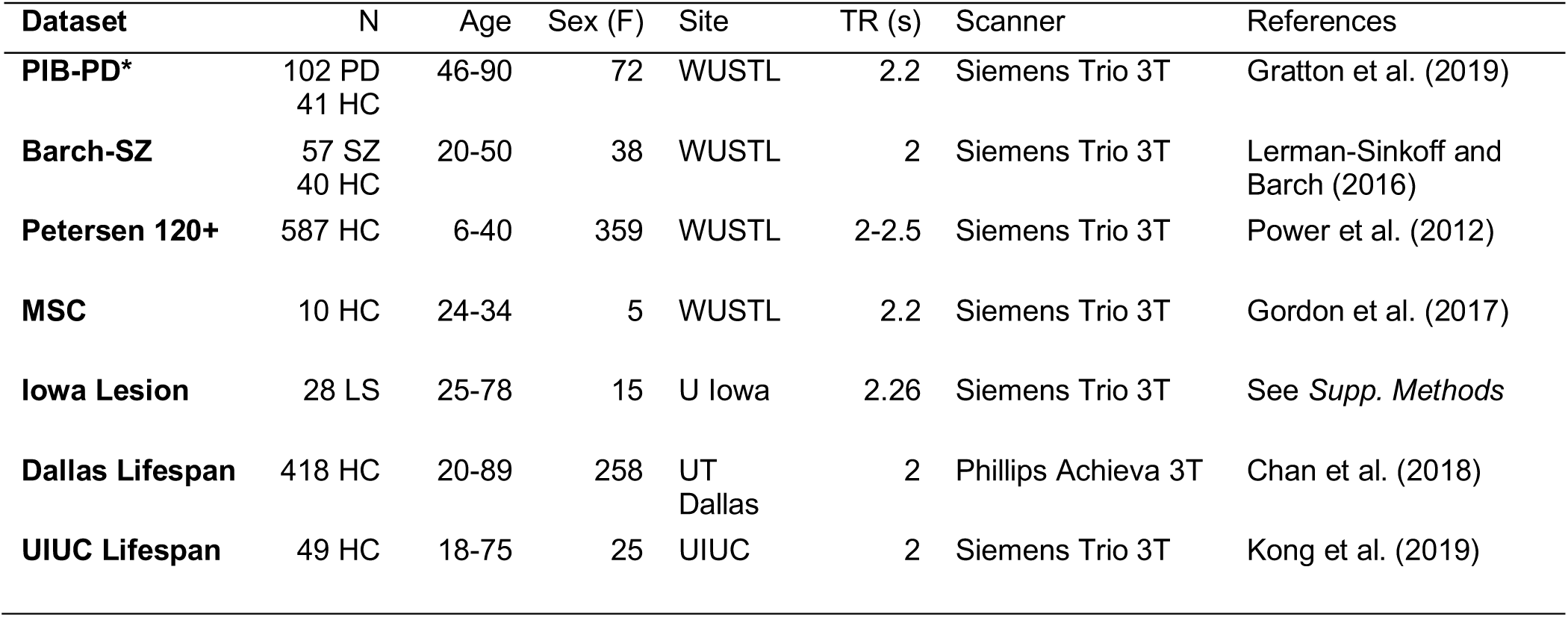
Dataset demographics and basic parameters. HC = Healthy control, PD = participant with Parkinson disease, SZ = participant with schizophrenia, LS = participant with brain lesion. WUSTL = Washington University in St. Louis, U Iowa = University of Iowa, UT Dallas = University of Texas at Dallas, UIUC = University of Illinois at Urbana-Champaign. *The PIB-PD Parkinson dataset is the primary dataset used in this paper for functional connectivity analyses.

| Dataset | N | Age | Sex (F) | Site | TR (s) | Scanner | References |
| --- | --- | --- | --- | --- | --- | --- | --- |
| **PIB-PD\*** | 102 PD<br>41 HC | 46-90 | 72 | WUSTL | 2.2 | Siemens Trio 3T | Gratton et al. (2019) |
| **Barch-SZ** | 57 SZ<br>40 HC | 20-50 | 38 | WUSTL | 2 | Siemens Trio 3T | Lerman-Sinkoff and Barch (2016) |
| **Petersen 120+** | 587 HC | 6-40 | 359 | WUSTL | 2-2.5 | Siemens Trio 3T | Power et al. (2012) |
| **MSC** | 10 HC | 24-34 | 5 | WUSTL | 2.2 | Siemens Trio 3T | Gordon et al. (2017) |
| **Iowa Lesion** | 28 LS | 25-78 | 15 | U Iowa | 2.26 | Siemens Trio 3T | See *Supp. Methods* |
| **Dallas Lifespan** | 418 HC | 20-89 | 258 | UT Dallas | 2 | Phillips Achieva 3T | Chan et al. (2018) |
| **UIUC Lifespan** | 49 HC | 18-75 | 25 | UIUC | 2 | Siemens Trio 3T | Kong et al. (2019) |

## Methods

### Overview

We analyzed data from several single-band scan sequences with repetition times (TRs) ranging from 2.0 – 2.5 seconds. Our goal was to characterize high frequency content in the realignment (motion) parameters and its effect on denoising for FC analysis. We first examined the relative prevalence of HF-motion. We then examined what participant characteristics are associated with HF-motion. Finally, we adjusted censoring-based denoising strategies to account for HF-motion via filtering and examined the consequences of this adjustment on FC. The approaches used in each of these steps are detailed below.

### Datasets

Data from seven different datasets were analyzed (**Table 1**). Four datasets were from Washington University in St. Louis (WUSTL), one dataset was from the University of Iowa (Iowa), one dataset was from the University of Texas at Dallas (Dallas), and one dataset was from the University of Illinois at Urbana-Champaign (UIUC). All datasets were collected with single-band scan sequences, with repetition times (TRs) ranging from 2.0 – 2.5 seconds. Collectively, these data, from both Siemens and Phillips 3T scanners, represent 1,332 participants (772 females), including children through older adults (ages 6 – 90), individuals with neurological and psychiatric disorders, and neurotypical controls. All procedures were approved by the Institutional Review Boards at the respective institutions.

The Protein and Imaging Biomarkers in Parkinson Disease (PIB-PD), Barch Schizophrenia (Barch-SZ), and Iowa Lesion datasets were included as they served as our initial observations of prominent HF-motion in single band datasets. The primary PIB-PD dataset was used to examine the consequences of HF-motion on functional connectivity. We added the Petersen 120+, Dallas Lifespan, and UIUC Lifespan datasets to examine the relationship between HF-motion and age, body mass index (BMI; Dallas, UIUC), and cardio-respiratory fitness (UIUC). The Midnight Scan Club (MSC) dataset was included to examine stability of HF-motion across repeated sessions.

**Table 1** includes a brief description of the characteristics of each dataset. Additional details on six of the seven datasets can be found in the associated references. Note that the Barch-SZ dataset only contains subjects from the second dataset (task runs) in Lerman-Sinkoff and Barch (2016). Details on participant characteristics and MRI data collection parameters in the final (Iowa Lesion) dataset can be found in the *Supp. Methods; Supp. Table 1* reports imaging parameters for all datasets. In all cases apart from the primary PIB-PD dataset, we included all participants who had a fully pre-processed dataset in our motion parameter analyses. The PIB-PD dataset had more restricted inclusion criteria as it was used for functional connectivity analysis, as described in the *Supp. Methods*.

### MRI collection parameters

Details about the MRI collection parameters for each dataset can be found in the respective references in **Table 1** and are summarized in **Supp. Table 1**. In brief, each dataset included fairly standard single band gradient echo EPI functional MRI scans on 3T scanners, with TRs ranging from 2 – 2.5s, with an A/P phase encoding direction. One dataset (UIUC Lifespan) was collected with GRAPPA (factor 2); no other datasets included GRAPPA, iPAT, or partial Fourier acceleration.

### Pre-processing and realignment

All functional data were rigid body realigned, thereby creating six rigid body motion parameter timeseries (3 translation and 3 rotation). This manuscript primarily focuses on analyses of the 6 rigid body parameters from this preprocessing. Rotation parameters were converted to mm, under the assumption of a 50mm head radius as in Power et al. (2012). When multiple runs were acquired in the same participant, the motion parameters were concatenated across runs before analysis. For the MSC dataset, where motion parameters were compared from sessions on different days, motion parameters from each session were kept separate. Realignment was conducted either using the 4dfp toolbox (https://sites.wustl.edu/nillabs/4dfp-documentation/; for the 4 WUSTL and the Iowa Lesion datasets), SPM8 (https://www.fil.ion.ucl.ac.uk/spm/software/spm8/; for the Dallas Lifespan dataset) or SPM12 (https://www.fil.ion.ucl.ac.uk/spm/software/spm12/; for the UIUC Lifespan dataset) using default parameters for rigid body realignment. Details on the analysis package, algorithm, and alignment reference frame are included in **Supp. Table 2**. All packages optimize the same objective function (the spatial correlation between each frame and the reference frame). Past work has demonstrated that FD traces are extremely similar (r≈0.95) regardless of reference volume, algorithm, or package (see Fig. S1 and associated analyses in (Power et al., 2017a)). Supplemental analyses in this manuscript also compare HF-motion estimates from different packages (4dfp; AFNI version 17.2.09; FSL version 5.0; see **Supp. Fig. 1**).

Slice timing correction was performed in all datasets before realignment. While this is a common strategy in analysis, the choice of slice timing correction influences FD metrics, as previously reported by Power et al. (2017a), and these effects are lessened with filtering (see **Supp. Figs. 2-3** for a more extensive illustration and discussion of the effects of slice timing correction on high-frequency motion). Field maps were only collected in a subset of datasets (MSC, Iowa Lesion) and when available, field map correction was applied to the functional data after realignment, and therefore will not influence the realignment parameters reported here.

### Analysis of motion parameters

#### Power spectral analyses

Power spectra of the motion parameters were calculated using the multitaper power spectral density (PMTM) module in Matlab. Power was then log-scaled and z-normalized within each motion parameter and expressed as a percentile, following (Fair et al., 2020). We refer to this scaled and normalized version of the power as “*<u>relative power</u>*”.

#### Quantification of high-frequency (HF) motion

HF-motion was quantified by determining the percent of relative power above 0.1 Hz within each motion direction. We focus on HF-motion in the y-translation (phase-encoding) direction given that factitious respiratory-related effects are largest in the phase-encoding direction (Brosch et al., 2002; Chen et al., 2019; Durand et al., 2001; Fair et al., 2020; Power et al., 2019; Raj et al., 2001; Van de Moortele et al., 2002). Supplemental analyses report the relationship between HF-motion in other directions and patient demographics.

#### Relationship between HF-motion and participant characteristics

The relative HF power in the phase-encoding direction was related to participant demographics including age and sex and, when available, diagnosis (PIB-PD, Barch-Sz), BMI (Dallas-Lifespan, UIUC-Lifespan), and cardiorespiratory fitness (UIUC-Lifespan; see measurement details in *Supp. Methods*). The relationship between HF-motion and participant demographics was tested within each dataset. Relationships between motion and individual demographics were examined using Spearman correlations (for continuous data) or two-sample two-sided *t*-tests (for sex, diagnosis). A final ANOVA was used to test the relationship between HF-motion and all variables at once within each dataset; participant demographics were z-scored before entry into the ANOVA. In each case *p*-values were corrected for multiple comparisons using FDR-correction across the number of tests (datasets); corrected p-values are reported as p(FDR). For some analyses, data from the WUSTL samples were grouped to create a broader age-range for analysis. Follow-up analyses used partial correlation to test for the role of each demographic variable after controlling for other demographic variables.

#### HF-motion filtering

To remove high-frequency content from the motion parameters, motion parameters were low-pass filtered at 0.1 Hz using a first order Butterworth filter with zero-padding (100 frames) on either end of the motion timeseries. The filter was applied both forward and backward and implemented with the *filtfilt.m* function in Matlab 2013b (see wrapper script *filter_motion.m* in the manuscript gitrepo https://github.com/GrattonLab). Low-pass filtering was conducted on all 6 motion parameters, not only the phase encoding direction, as cross-talk may occur between realignment parameters (see Appendix A in Fair et al. (2020)).

#### Framewise Displacement (FD) vs. filtered Framewise Displacement (fFD)

As in Power et al. (2012), FD represents the sum of the absolute frame-to-frame difference in motion parameters. Framewise displacement was calculated on both the original motion parameters (*<u>FD</u>*) as well as the low-pass filtered motion parameters (*<u>fFD</u>*).

### Functional Connectivity (FC)

#### FC Analysis

In the PD dataset, additional processing steps were done to align and denoise the functional and structural MRI data for FC analyses. These steps are detailed in the *Supplemental Methods*. Afterwards, denoised FC was related to motion measures (see below) to assess the utility of FD vs. fFD metrics in censoring for removing motion bias in FC.

#### FD vs. fFD based censoring

High motion frames were identified using either the FD or fFD metric. With FD, frames were marked as “high motion” if they had values above 0.2 mm (as in (Power et al., 2014a)). The fFD measure requires lowering the censoring threshold as filtering reduces the amplitude of the FD measure (a natural consequence of applying a filter). Two different criteria were tested: a more lenient 0.1 mm threshold and a more conservative 0.08 mm threshold. For functional connectivity analyses, censoring masks also included removal of 14 frames at the start of each run (Laumann et al., 2015) and short frame segments (<5 long) (Power et al., 2014a). For the primary versions of these analyses, each participant was required to have 50 low-motion frames per run and 150 low-motion frames total. Different total and segment minima were also compared in supplemental analyses.

#### Motion-FC (QC-FC) Analysis

Additional analyses were conducted following Ciric et al. (2017) to test for the relationship between motion and functional connectivity measures (i.e., QC-FC analyses). These analyses correlated mean framewise displacement (QC) with functional connectivity for a given connection across participants. Given that censoring approaches most strongly influence the distance-dependence of QC-FC relationships (Ciric et al., 2017; Parkes et al., 2018; Power et al., 2012; Satterthwaite et al., 2012), we focus primarily on these analyses to evaluate different censoring strategies. The motion QC measure adopted in most analyses was based on the mean of the fFD measure, but supplemental analyses also report QC based on mean FD. Additional analyses adapted from Ciric et al. (2017) of the median QC-FC relationship, the percent of significant QC-FC relationships, and the number of QC-FC relationships above r=0.4 are also reported in the supplement.

Statistical evaluation of QC-FC distance dependent slopes of each censoring strategy was conducted by comparison with a null random censoring approach. For a given subject, random censoring involved removal of the same number of frames as called for in the true censoring strategy, but in this case censored frames were randomly selected rather than selected for their high FD (or fFD) values. Frame matching was done per participant. N=100 permutations of random frame censoring were run; in each permutation, the same FD-FC benchmark tests were conducted and stored. Thus, random censoring represents a null baseline comparison which is matched for number of frames.

### Data & Code Availability

Code associated with this manuscript will be made available upon publication at https://github.com/GrattonLab. Data from the Midnight Scan Club is available at https://openneuro.org/datasets/ds000224. Data associated with the first 120 subjects from the Petersen 120+ dataset are available at https://openneuro.org/datasets/ds000243/versions/00001. Other datasets are available upon request.

## Results

### High frequency motion is present in single-band fMRI data, especially in the phase-encoding direction

The present investigation was motivated by observation of a phenomenon in the realignment parameters of two datasets that included older participants (PIB-PD and Iowa Lesion) that was unexpected to us, and appears relatively underappreciated in the field. To demonstrate the characteristics of this observation, we compare realignment parameters and their spectra from two participants in the PIB-PD dataset (**Fig. 1A**; the PIB-PD dataset will be used as our primary dataset throughout this manuscript). One of these participants (**Fig. 1A top**) shows typical realignment parameters for someone with low head movements (97% of frames are below an FD threshold of 0.2 mm). The second participant (**Fig. 1A bottom**), instead, has a prominent zig-zag pattern in the realignment time-courses most clearly in the y-translation (phase-encoding) direction. This is also seen in the spectra of the realignment parameters, where a large peak in the relative power of the y-direction realignment parameter is seen above 0.1 Hz which we term *high-frequency motion* (*HF-motion*). Besides this HF-motion, the second participant also appears to show low overall movement, much like the first. However, because of the HF-motion, only 31% of frames are below an FD threshold of 0.2 mm. **Supp. Figure 1** shows that similar results are found when the motion parameters from these two participants are derived from different algorithms (AFNI’s 3dvolreg, FSL’s mcflirt), suggesting that HF-motion is not due to a particular analysis stream.

**Figure 1:**
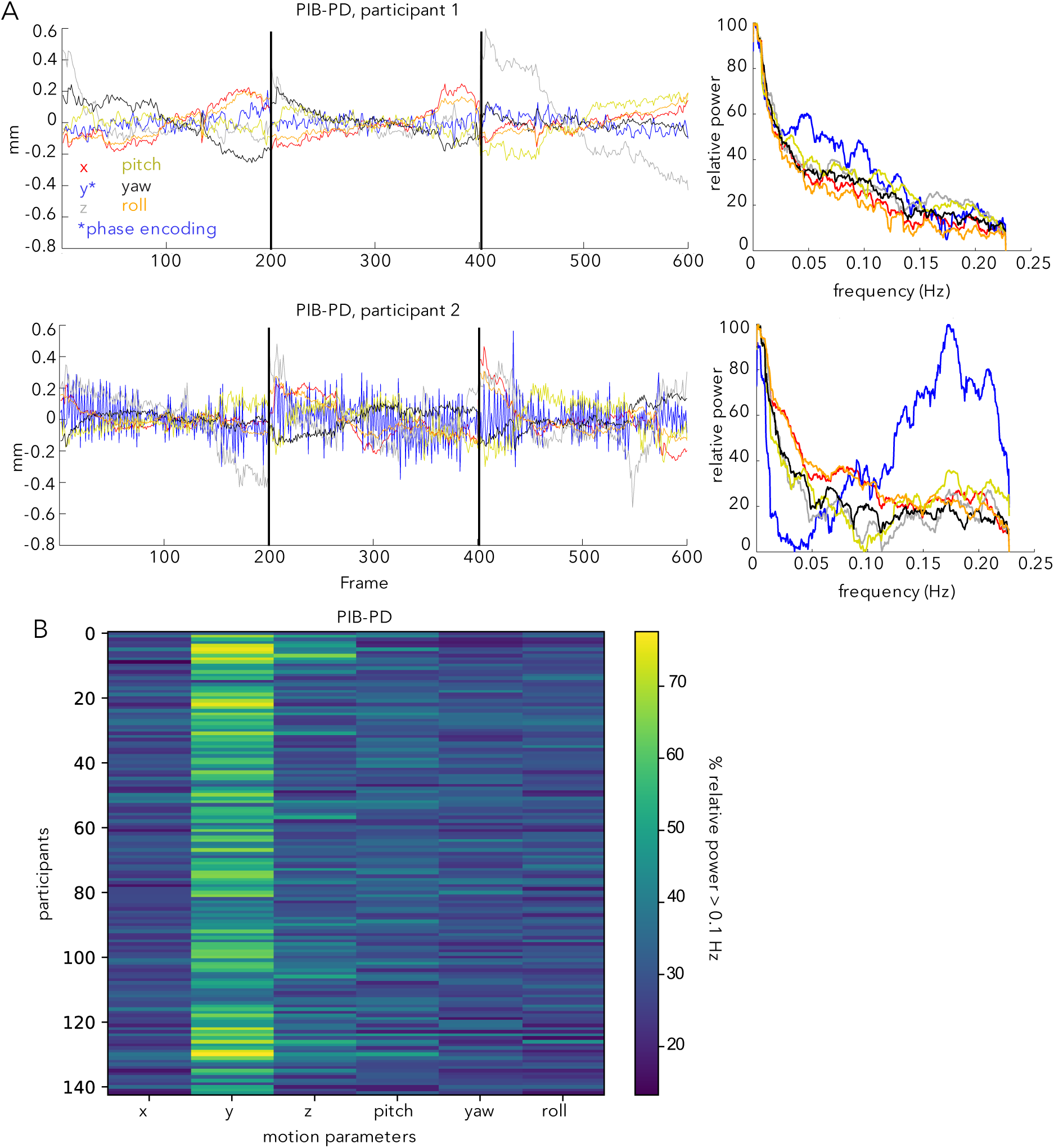
(A) Motion parameters in the PIB-PD dataset. Two participants are illustrated, one with little HF-motion (top) and one with substantial HF-motion (bottom). Left side plots depict motion parameters over time; right side plots depict the power spectra of the motion parameters expressed in terms of relative power (see *Methods*). The bottom participant shows a predominance of power above 0.1 Hz in the y-translation (\**phase-encoding*) direction (blue line). Black lines mark run boundaries. (B) Plot depicting the percent of relative power above 0.1 Hz. in each of the six motion parameters across all (N=143) PIB-PD participants. Y-translation shows the most HF-motion.

Across the full set of participants in the PIB-PD dataset (**Fig. 1B**, see also **Fig. 2A**), many of the PIB-PD participants show some evidence for this HF-motion, which is strongest in the phase encoding (y-translation) direction (**Fig. 1B**). HF-motion appears next most frequently in the z-direction but with substantially reduced magnitude (**Fig. 1B**; paired t-test comparing percent of relative HF-motion in y-translation and z-translation directions: t(142) = 18.32, p<0.001). The correlation between HF-motion in the y- and z-directions was modest, at r = 0.29. Across participants, HF-motion varies somewhat in peak frequency and magnitude (**Fig. 2A**).

**Figure 2:**
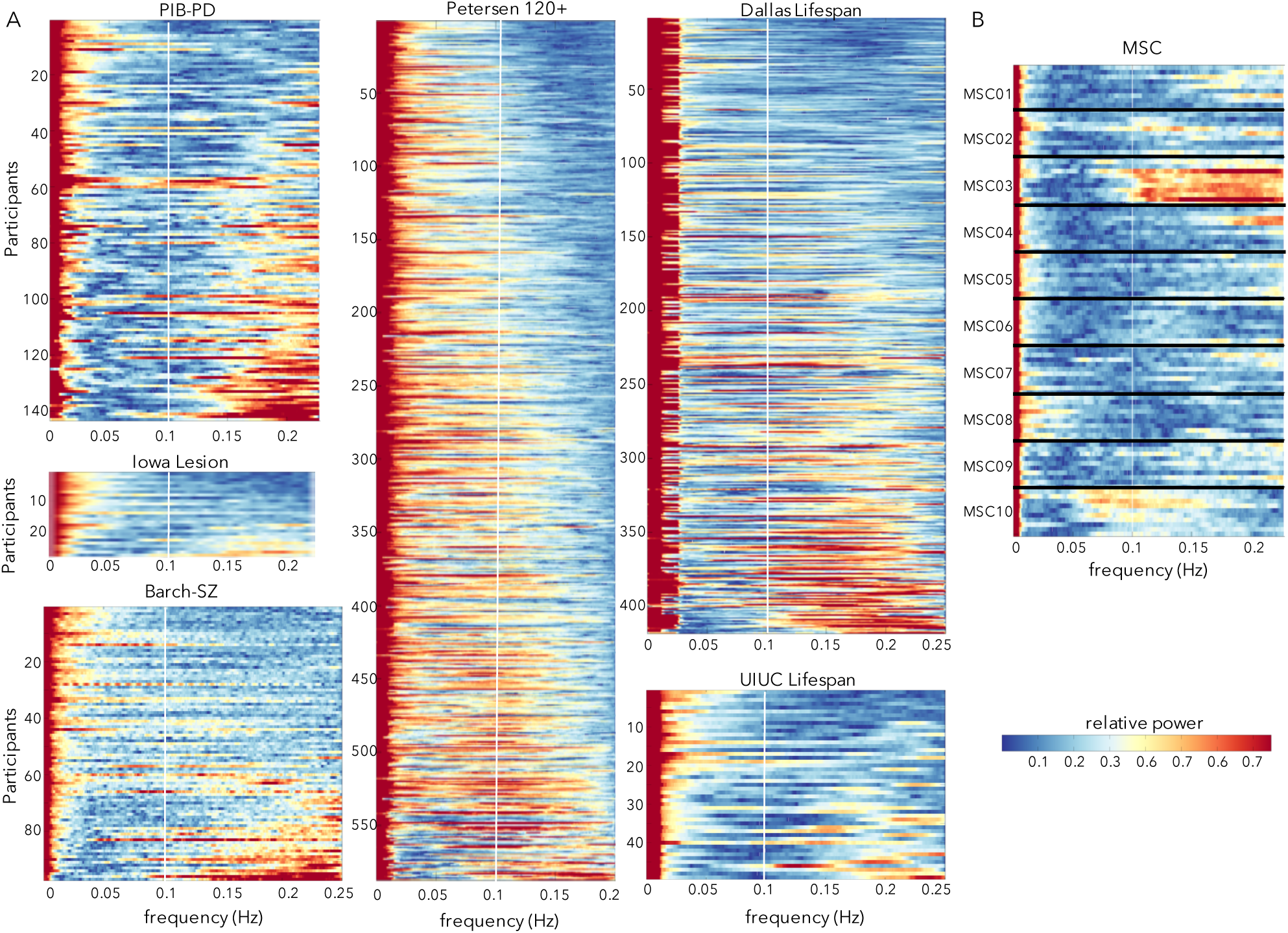
Power spectra of the y-translation (phase-encoding) motion parameter across diverse datasets. (A) Power spectra in six datasets with TR from 2 – 2.5 collected at multiple sites and in different populations. Participants are ordered based on their proportion of HF-motion. The white line marks 0.1 Hz; at these sampling frequencies respiratory effects in adults would typically alias into the 0.1 – 0.2 Hz range (above the white line). A set of participants in each dataset show peaks in power above 0.1 Hz. (B) Power spectra from 10 sessions in the 10 MSC subjects. For a given participant, each row depicts a single scan session and subjects are separated by horizontal black lines. HF-motion shows some consistency across sessions, with participants who exhibit the strongest HF-motion (e.g., MSC03) showing strong HF-motion in most repeated sessions (e.g., note similarity in frequency profile of motion parameters across repeated scans – stacked lines – from the same subject; MSC03 has 8/10 sessions with >67% of power in the y-translation above 0.1 Hz; other subjects show 0-1 sessions).

HF-motion is evident also in several other datasets, including those collected on the same scanner at WUSTL (Barch-Sz, Petersen 120+) and those collected at other sites (Iowa Lesion, UIUC Lifespan) and from a different scanner manufacturer (Dallas Lifespan – Phillips Achieva), all with TRs ranging from 2 – 2.5 s (**Fig. 2A**). In all datasets, a subset of individuals shows prominent HF-motion above 0.1 Hz, with relative power above 0.5, indicating greater than 50^th^ percentile HF power levels.

In datasets with repeated scans from the same individual (Gordon et al., 2017), HF-motion characteristics show stability over sessions (**Fig. 2B**): that is, if an individual shows HF-motion in one session, they are likely to show it again in other sessions at similar frequency ranges. In particular, MSC03 from MSC dataset shows consistent HF-motion above 0.1 Hz. MSC10 also shows a consistent peak in motion estimates, though centered on a lower frequency (around 0.1 Hz). Notably, in the MSC dataset, participants who exhibited the most drowsiness and prominent low frequency motion (MSC08) were different from those that had the most HF-motion (MSC03, and to a lesser extent MSC10 and MSC01).

Thus, HF-motion is relatively commonplace, although more prevalent in some participants and datasets (PIB-PD, Barch-Sz) than others (Petersen 120+, MSC).

### High frequency motion is related to age, BMI, and cardiorespiratory fitness

We next investigated whether the presence of HF-motion is linked to participant characteristics. Past work suggests links between HF-motion in the phase-encoding direction and main field perturbations caused by chest-wall motion (Brosch et al., 2002; Fair et al., 2020; Power et al., 2019; Raj et al., 2001). Thus, we examined participant characteristics potentially related to this phenomenon, i.e., age, sex, body mass, and cardiorespiratory fitness, which may modulate both respiratory rates and chest size.

Since we first noted HF-motion in datasets with older participants (PIB-PD, Barch-Sz, Iowa-Lesion), we began by examining whether HF-motion (defined as the proportion of relative motion > 0.1 Hz in the phase-encoding direction) varies across datasets sampling different age ranges. Consistent with our impression, we found significantly stronger HF-motion in datasets with older participants relative to the younger Petersen-120+ dataset (Petersen vs. PIB-PD: *t*(929) = 18.14, *p* = 3.27*10^−63^, *p*(FDR)<0.001; Petersen vs. Barch-Sz: t(883) = 17.84, *p* = 4.86*10^−61^, *p*(FDR)<0.001; Petersen vs. Iowa-Lesion: t(814) = 2.35, *p*=0.0191 p(FDR) = 0.0196).

By combining the PIB-PD, Barch-Sz, and Petersen-120+ datasets, we were able to examine the correlation between age and HF-motion in participants from 6-90 years old scanned on the same Siemens 3T Trio scanner at WUSTL (**Figure 3A)**. We found a significant relationship between HF-motion and age (Spearman rank correlation: ρ = 0.53, *p* = 8.35*10^−60^, p(FDR)<0.0001; if restricted to control participants only: ρ = 0.44, *p* = 1.92*10^−29^). This correlation is primarily driven by a rise in HF-motion across the young to middle aged participants in the Petersen 120+ group (ρ = 0.21, p = 2.99*10^−7^, p(FDR)<0.0001). To further confirm this finding, we obtained data from two adult lifespan fMRI datasets with participants from 20 through 90 years old (Dallas Lifespan and UIUC Lifespan; **Fig. 3B**). A significant relationship to age was found in the Dallas Lifespan dataset (ρ = 0.26, p = 1.15*10^−7^, *p*(FDR)<0.0001; note similar magnitude to the Petersen 120+ finding). No relationship was seen in the smaller UIUC Lifespan dataset (ρ = 0.03, *p* = 0.812). We speculate this may be due to the small size and unusual characteristics of the oldest participants in this study which required attendance of three sessions for fMRI, optical imaging, and behavioral measurements (note the downward trend from 70+). To further investigate whether age was related to HF-motion in a non-linear fashion, we used a smoothing spline approach to relate age and HF-motion in each of the three datasets (**Supp. Fig. 4**). With this approach, all three datasets show evidence of upward trends in early to middle age ranges, but a downward trend at the oldest ages. However, the exact shape of these trends differed across datasets (both the Dallas Lifespan and WUSTL datasets suggest a decline after 75; the UIUC Lifespan dataset shows a decline starting earlier around age 60, but has relatively sparse sampling of these ages). Thus, age is related to HF-motion in most datasets, but may exhibit a non-linear relationship with age that depends on the demand characteristics of each study.

**Figure 3:**
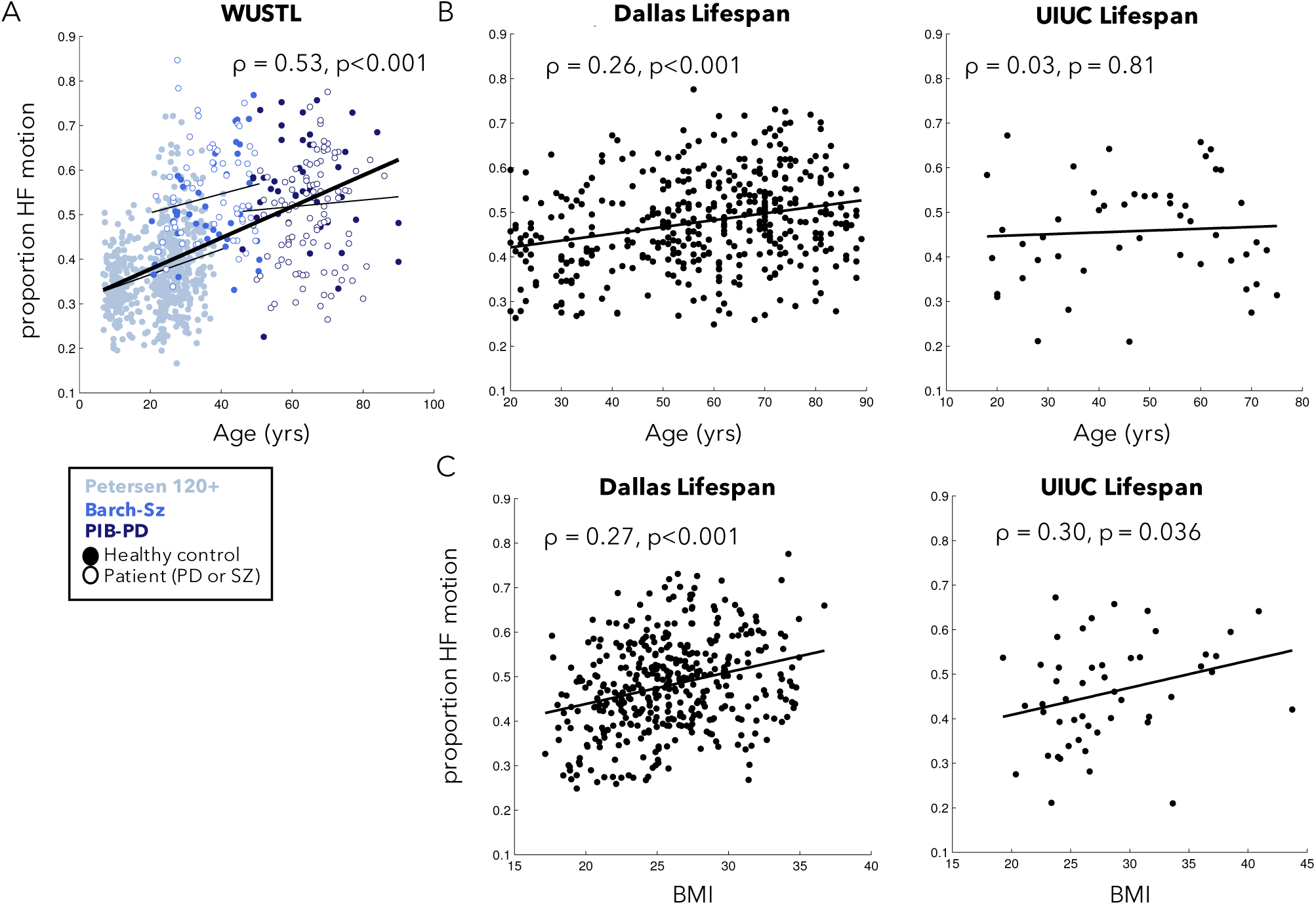
Relationship between phase-encoding HF-motion and participant age and BMI. (A) HF-motion showed a significant relationship to age in a combined set of three datasets collected on the same scanner at WUSTL. Different colors represent data from different datasets, with the fit line plotted for each dataset separately as a thinner black line. Filled circles represent data from healthy controls and open circles represent data from individuals with Parkinson Disease (PD) or schizophrenia (SZ). (B) HF-motion was also related to age in the Dallas lifespan dataset, but not in the smaller UIUC lifespan dataset. (C) HF-motion was related to participant BMI in both the Dallas and UIUC Lifespan datasets. ***Supp. Fig. 6*** shows scatter plots after correction for alternate participant characteristics.

Next, we examined whether HF-motion relates to participant BMI given the potential connection between HF-motion and chest-wall deviations (**Figure 3C**). BMI measures were only collected in the Dallas Lifespan and UIUC Lifespan datasets. We find a significant and similar relationship between HF-motion and BMI in these two datasets (Dallas Lifespan: ρ = 0.27, p = 4.85*10^−8^, p(FDR)<0.0001; UIUC Lifespan: ρ = 0.30, p = 0.0364, p(FDR) = 0.0394). The UIUC Lifespan dataset also collected measures of cardiorespiratory fitness (see *Supp. Methods* for description) and, interestingly, this measure also showed a modest relationship to HF-motion (ρ = −0.31, p = 0.030; **Supp. Fig. 5**). In the Dallas Lifespan dataset the relationship of HF-motion with BMI and age remain significant when controlling for the alternate variable via partial correlation (HF-motion vs. BMI, controlling for age: ρ = 0.26, p = 9.99*10^−8^; HF-motion vs. age, controlling for BMI: ρ = 0.25, p = 5.61*10^−7^; **Supp. Fig. 6**). In the UIUC Lifespan dataset, the relationship between HF-motion and cardiorespiratory fitness remained significant after controlling for age and BMI (ρ = −0.34, p = 0.02), but the relationship between HF-motion and BMI is somewhat weakened and no longer significant after controlling for age and cardiorespiratory fitness (ρ = 0.23, p = 0.12; **Supp. Fig. 6**). This may be due to the smaller number of participants in the UIUC Lifespan dataset and/or to the association between BMI and cardiorespiratory fitness (cardiorespiratory fitness vs. BMI: r = −0.37, p<0.01; Age vs. BMI: r = 0.11, p = n.s.). Generally, these findings suggest that older, higher BMI and/or less fit participants may be more likely to show HF-motion. For completeness, we also examined the relationship between HF-motion in other motion directions and these demographic variables (**Supp. Fig. 7**); the findings above were to some extent shared across motion directions, with the y-translation (phase-encoding) direction showing the highest relationship on average, followed by the z-translation. Shared relationships may be driven by cross-talk between realignment parameters (see Appendix A in (Fair et al., 2020)).

Additional differences in HF-motion based on gender (females > males in the Petersen 120+) and diagnosis (PD participants < HC controls, SZ participants > HC controls) were also observed but less consistent across datasets (**Supp. Fig. 5**). A full table of ANOVA results relating participant characteristics to HF-motion is included in **Supp. Table 3**. These results suggest the potential concern that HF-motion may affect group comparisons unevenly.

### High frequency motion disrupts the relationship between FD and BOLD signal abnormalities

In the previous sections we demonstrated the presence of HF-motion in several datasets, as well as a link to participant characteristics. In this and the next few sections we examine how HF-motion influences FC analyses.

Over the past decade, numerous studies have demonstrated a relationship between head movements (even small, ∼0.2mm) and BOLD signal artifacts that bias functional connectivity (Power et al., 2012; Satterthwaite et al., 2019; Satterthwaite et al., 2012; Van Dijk et al., 2012) (e.g., see **Fig. 4A**). Typically, in these studies, head motion is summed across the 6 realignment parameters to calculate Framewise Displacement (FD). Methodological studies suggest that removal of frames with elevated FD (frame censoring) can effectively reduce bias in functional connectivity (Ciric et al., 2017; Power et al., 2014a; Satterthwaite et al., 2019; Satterthwaite et al., 2012; Satterthwaite et al., 2013). However, these methodological studies primarily focused on the analysis of child or young adult data which, as reported in the previous section, exhibit significantly less HF-motion than datasets acquired in middle aged or older participants.

**Figure 4:**
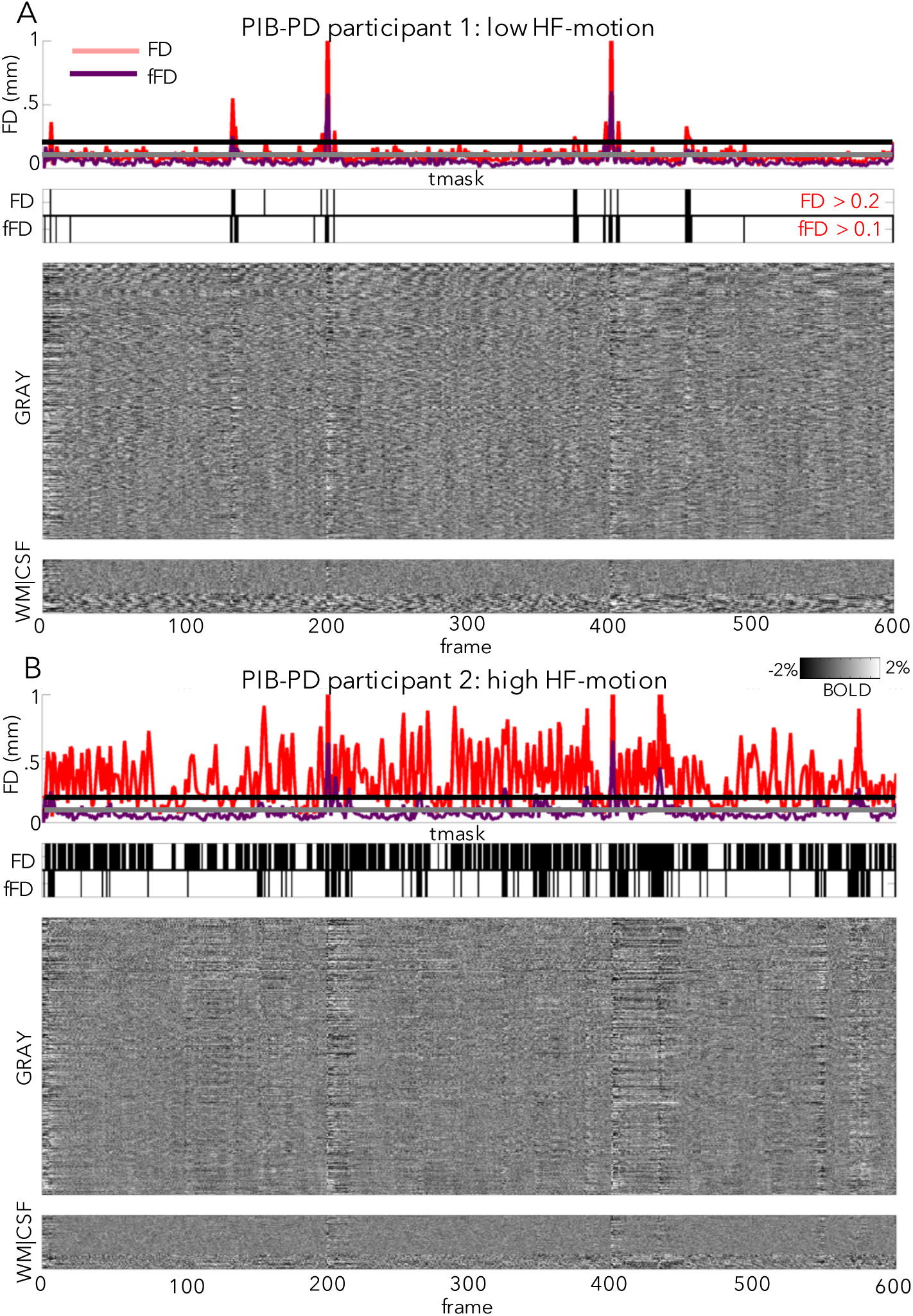
Relationship between fMRI signal changes and FD. Here we plot the relationship between FD (a proxy for motion) and fMRI signal changes (via fMRI intensity “grayplots” (Power, 2017); see *Supp. Methods*) in a participant with low HF-motion (A) and a participant with high HF-motion (B) in the PIB-PD dataset. In each panel, the top subplot shows the FD trace (red = standard FD, purple = FD calculated from filtered motion parameters, horizontal lines marking typical thresholds for censoring: black = 0.2 for FD and gray = 0.1 for filtered FD). The second subplot shows a mask of frames above threshold that will be marked for censoring (top row: FD, bottom row: filtered FD). The third subplot shows the stacked fMRI timecourses for gray matter regions (scale: −2:2% signal change). The final subplot shows stacked timecourses for white matter and CSF regions. As can be seen, censoring masks are quite different in in subjects with large amounts of HF-motion, but relatively similar in subjects with low HF-motion. *NB: this grayplot is presented after pre-processing, demeaning and detrending, and nuisance regression, but before interpolation and temporal filtering; frame censoring can affect multiple stages of processing, including application of nuisance regression if only “low motion” frames are used for nuisance fits (Power et al., 2014a)*.

In our older adult datasets, participants with low HF-motion (**Fig. 4A**), show the characteristic relationship between FD spikes and BOLD signal artifacts (e.g., note correspondence of vertical stripes in the BOLD signal with raised FD). However, participants with prominent HF-motion show a disrupted relationship between FD measures and BOLD signal artifacts, driven by a raised baseline of the FD timecourse (**Fig. 4B**). This is expected, as HF-motion will strongly drive frame-to-frame FD. Thus, a very high proportion of frames are flagged as having elevated FD (FD > 0.2, (Power et al., 2014a)) and marked for censoring in these participants, even in the absence of discernable artifact in the BOLD signal (e.g., note the absence of vertical bands at most of these time-windows). Thus, with HF-motion, standard FD appears to become a poor marker of true micro-movements and BOLD signal disruptions.

### Filtering motion parameters before FD calculation as an approach to removing HF confounds

One possibility is that FD values contaminated by HF-motion may be improved by filtering motion parameters before FD calculation to remove the HF-motion component. We tested this hypothesis, by low-pass filtering motion parameters at 0.1 Hz. We then used these filtered parameters to create a new *filtered FD* (or *fFD*) measure. These filtered motion traces closely resemble the original motion traces, still showing evidence of transient small movements, and indeed are almost identical in participants with low HF-motion. However, in participants with high HF-motion, the prominent high frequency flickering is removed (see **Supp. Fig. 8** for examples of filtered motion parameters from the same subjects shown in **Fig. 1**).

Because we remove power from the motion parameters, fFD tends to show a lower baseline relative to the original FD measure, even in low HF-motion subjects (see **Fig. 4A**, compare FD to fFD line in top subplot). Thus, it is necessary to lower the FD threshold for censoring to more effectively capture small head movements (Power et al., 2015). In this and the next section, we tested the results of censoring motion frames based on a filtered FD threshold of 0.1 mm. or a more conservative 0.08 mm (see **Supp. Fig. 9-10** for justification for selecting these values).

In participants without HF-motion (**Fig. 4A**), FD and fFD appear to be quite similar. In subjects with HF-motion, the fFD measure appears to better align with fMRI signal artifact and more effectively expose head movements (**Fig. 4B**). Moreover, despite using a more conservative threshold, fFD leads to a much higher proportion of frames retained in these subjects (**Fig. 4B**, compare censoring masks^2^). These initial observations are quantified in the next section.

### Filtering motion parameters before FD calculation saves data from frame censoring

Next we quantified these effects across all participants in the PIB-PD dataset. As the examples suggested, using fFD saves a substantial amount of data from censoring in many participants, despite the more conservative threshold (**Fig. 5A**, points above the unity line). On average, 15.7% fewer frames were flagged when fFD<0.1 was used for censoring relative to FD<0.2 (std = 16.6% of frames; t(142) = 11.29, p<0.001). Moreover, if we require 150 low motion frames to include a participant in final analyses (∼5.5 min.), then fFD<0.1 censoring leads to retaining an additional 11.9% of participants in the PIB-PD dataset, relative to FD<0.2. Smaller but still significant data savings were seen with the more conservative fFD<0.08 threshold (**Supp. Fig. 11**; average frame savings: 6.4%, std = 16.6%, t(142) = 4.61, p<0.001; 7.7% more subjects retained). Similar results were obtained in the other datasets with large amounts of HF-motion (**Table 2, Supp. Fig. 12**; note the exception of the younger Petersen 120+ dataset).

**Table 2:**
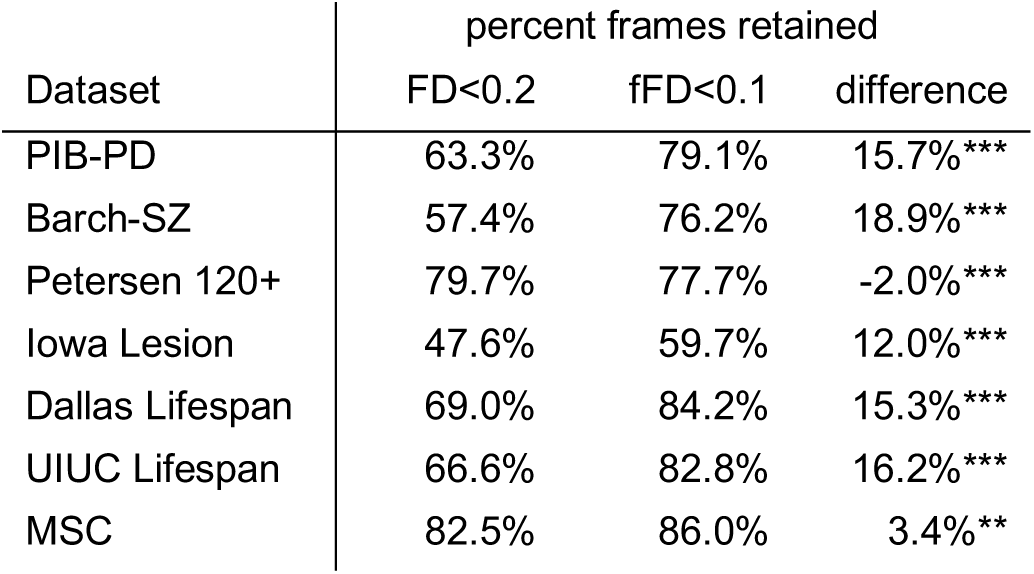
Comparison of frames retained with FD<0.2 or fFD<0.1 criteria in different datasets. Across most datasets, adopting fFD<0.1 for frame censoring retained significantly more frames that an FD<0.2 criteria. The exception is the Petersen 120+ dataset. ***p<0.001, **p<0.01

**Figure 5:**
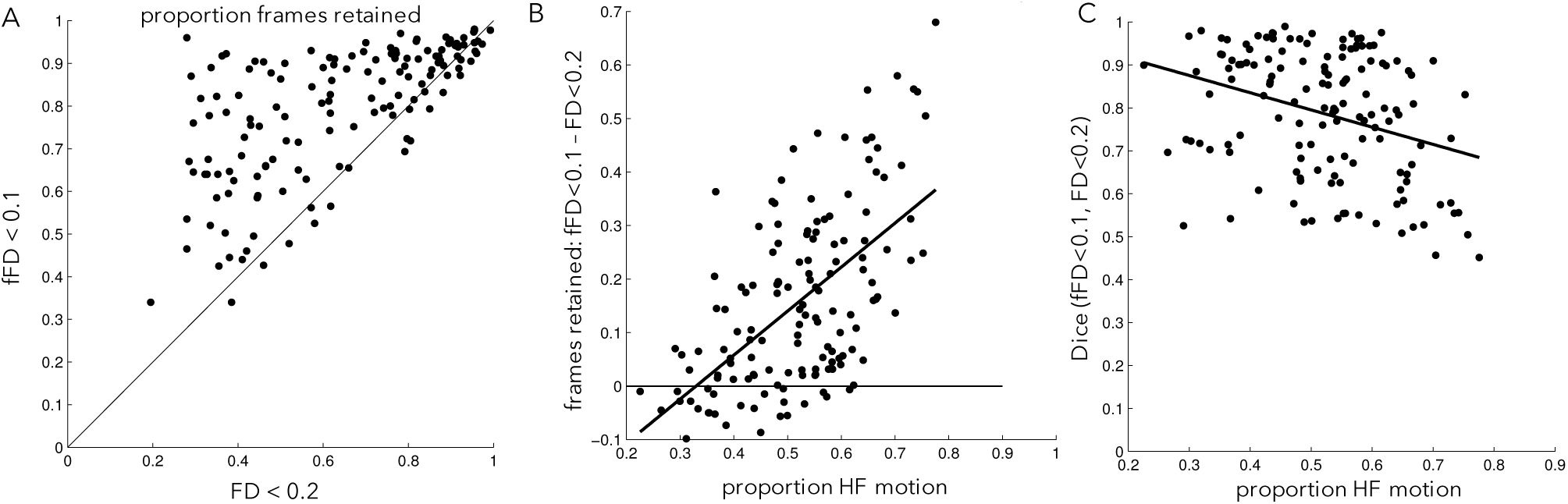
FD vs. fFD relationship in the PIB-PD dataset. (A) Proportion of frames below an FD threshold of 0.2 (x-axis) or an fFD threshold of 0.1 (y-axis). Black line is unity. Note that most points lie above the black line, indicating data savings despite a lowered fFD threshold. (B) The differences in frames retained between the FD<0.2 and fFD<0.1 is related to the amount of HF-motion (y-direction) in a given participant. (C) The similarity between censoring masks created from the fFD<0.1 threshold and the FD<0.2 threshold is related to the amount of HF-motion in a given participant. Participants exhibiting low levels of HF-motion show the most similar censoring masks.

As might be expected, data savings were largest in participants with more HF-motion (**Fig. 5B**, ρ = 0.55). In general, censoring masks constructed using filtered and unfiltered FD had high overlap, but the similarity between the two masks was correlated with the amount of HF-motion: they looked most similar in participants with little HF-motion (**Fig. 5C**; ρ = −0.30, p<0.001).

In summary, these results demonstrate that filtering motion parameters before calculating FD can save substantial amounts of data, particularly in participants with HF-motion contamination. Concomitantly, those with little HF-motion artifact are not particularly affected by filtering and show very similar censoring masks regardless of the approach.

### Filtering motion parameters before FD calculation does not bias functional connectivity measures

Filtering motion parameters before FD calculation saves data from censoring. But, does it still address motion confounds in fMRI functional connectivity analyses? The stronger link between fFD and fMRI artifacts (**Fig. 4**) suggests that it might; here we more formally test this question by applying the benchmark criteria from Ciric et al. (2017) to our processing stream with censoring using either FD or fFD criteria.

Past work (Ciric et al., 2017; Parkes et al., 2018; Power et al., 2012; Power et al., 2014a; Satterthwaite et al., 2012) suggests that frame censoring techniques exert the strongest influence on distance-dependent biases in functional connectivity. Without frame censoring, motion biases functional connectivity based on the distance between regions: when regions are close together, functional connectivity is relatively amplified and correlated with motion, whereas when regions are far apart, functional connectivity is relatively dampened. Frame censoring addresses this confound, reducing the link between connection distance and motion. Accordingly, here we focus first on whether the filtered FD censoring criterion still adequately addresses distance-dependent bias in functional connectivity data, at a similar level to that previously observed for FD criteria in the absence of prominent HF-motion.

The following analysis is restricted to a set of N = 79 PIB-PD participants (52 PD and 27 HC) who had >150 frames under all three frame censoring criteria (FD<0.2, fFD<0.1, fFD<0.08) to ensure that observed differences could be attributed to censoring rather than subject inclusion. Note that from here onward, all censoring masks also include removal of 14 frames at the start of each scan as well as censoring of small segments less than 5 frames long (Power et al., 2014a) unless otherwise noted. **Supp. Table 4** summarizes the mean and range of frames retained under each censoring strategy within this restricted sample.

Distance-dependent relationships between FC and motion were small in this dataset using any of the three censoring criteria (**Fig. 6A**, compare with a QC-FC distance-dependence of r = −0.116 in (Ciric et al., 2017)), all of which did better at eliminating bias than analysis strategies omitting censoring (**Fig. 6B**). Importantly, distance-dependent bias was not elevated when using fFD for censoring rather than FD, despite the inclusion of a large number of additional frames. All three censoring strategies out-performed random censoring, with z-statistics relative to their matched null distributions of z=5.14 for FD<0.2, z=8.43 for fFD<0.1, and z=9.76 for fFD<0.08. With N=100 random permutations, these values are all significant at p<0.01. See **Supp. Fig. 13** for different variations on this analysis, varying the quality metric, contiguous segment length, and minimum number of frames per participant. We note that inclusion of fewer frames per participant tended to increase distance-dependent bias (**Supp Fig. 13C**). Regardless of the specific implementation, we consistently find that the three censoring strategies perform similarly, better than the non-censoring based approaches, and better than random-censoring based approaches. Thus, we conclude that use of fFD censoring approaches can save data substantial data while still adequately addressing motion confounds.

**Figure 6:**
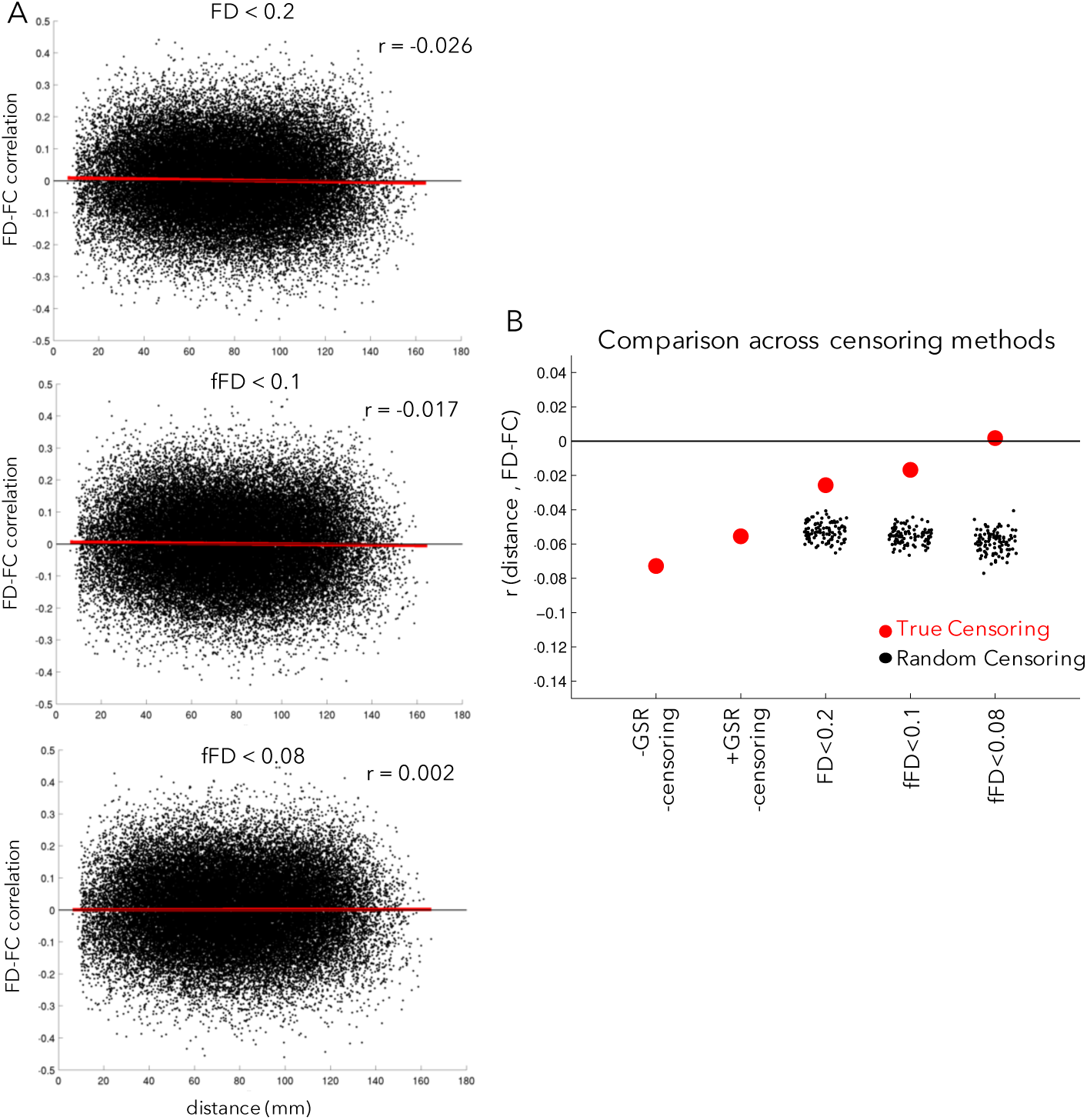
Relationship between motion and distance in FC. (A) The x-axis plots the distance between a pair of regions and the y-axis plots the correlation between the FC of those regions and motion (mean fFD) across participants for each pair of regions (black points), as in (Ciric et al., 2017). The red line shows the linear fit. In the absence of correction, head motion typically leads to elevated FD-FC relationships for regions close together in space and relatively dampened FD-FC relationships for regions far apart, leading to an overall negative slope. In the top plot, censoring has been conducted using an FD < 0.2 criteria, in the middle plot using a filtered FD < 0.1 criteria, and in the bottom plot using a filtered FD < 0.08 criteria. (B) Summary of the average distance-dependent correlations across censoring strategies (red dots); for comparison, we also computed distance-dependent relationships with random censoring (black dots). Filtered FD censoring criteria do well at removing distance-dependent bias in functional connectivity (relative to FD<0.2 censoring, random censoring, or absent censoring) while retaining substantially more data. For variations on these analyses, see **Supp. Fig. 13**.

We next sought to replicate these findings in another dataset, electing to use the frequently studied (Power et al., 2012; Power et al., 2014a) and accessible Petersen 120+. We restricted our analyses to 100 subjects in the Petersen 120+ with the highest levels of HF-motion (based on the relative HF-motion metric). Of these subjects, N = 81 passed our criteria for minimum data quantities (>150 frames) for functional connectivity analysis under both fFD<0.1 and FD<0.2 censoring approaches. The results of this comparison are shown in **Supp. Figure 14**. As with the PIB-PD dataset, both FD<0.2 and fFD<0.1 do similarly at reducing QC-FC distance dependence, well out-performing random censoring of the timeseries (with N=100 permutations, p<0.01). Thus, the good performance of filtered FD parameters at reducing distance-dependent artifact replicates in another independent dataset (notably sampling a very different population).

In addition to distance-dependent bias, we examined other benchmarks for adequate motion artifact removal from Ciric et al. (2017) in the PIB-PD sample (**Supp. Fig. 15**). Again, fFD criteria did well, at levels similar to FD based censoring. In these cases, frame censoring had a relatively small effect as compared to global signal regression, as has previously been reported (Ciric et al., 2017; Parkes et al., 2018). As might be expected, given these findings, group correlation matrices in PD and HC groups are quite similar across these three censoring strategies (**Supp Fig. 16**). Thus, these results indicate that filtering motion parameters before FD calculation saves data without re-introducing biases in functional connectivity analysis.

## Discussion

We found that high frequency fluctuations in motion (HF-motion) that have previously been related to respiration in fast TR fMRI datasets are also present (in aliased form) in common single band TR datasets. We demonstrate that HF-motion is evident across seven datasets representing diverse populations, scanner protocols, and sites. HF-motion is more prevalent in older participants, those with higher BMI, and those with lower cardiorespiratory fitness, suggesting that it may pose a larger issue in populations with these characteristics. We show that the presence of HF-motion reduces the link between motion metrics and fMRI signal quality and suggest that filtering motion parameters before FD censoring may be an effective strategy for correcting this contamination. We demonstrate that this filtering approach saves a substantial amount of data from censoring in most datasets, especially those acquired in middle-age to older adults (12-19% of data savings in these samples). Furthermore, we show that filtered motion parameters appear to demonstrate greater fidelity to artifacts in fMRI data. Functional connectivity benchmarking analyses demonstrate that filtered motion parameters are still effective at reducing motion biases in functional connectivity. These results promise substantial data savings in many common analysis pipelines that incorporate frame censoring.

### Sources of HF-motion in realignment parameters: factitious and true head motion

Fair et al. (2020) recently demonstrated the presence of high-frequency fluctuations in the motion parameters of a multiband dataset at frequencies that closely match respiration rates. This finding was corroborated by several additional reports of high frequency “motion” fluctuations in other multiband datasets (Chen et al., 2019; Etzel, 2016a; Inglis, 2016b; Power et al., 2019). At least two hypothetical mechanisms may link respiration with changes in realignment parameters. One is that true head movements may be induced by the physical connection between the head and chest during respiration. The second is that respiration can induce factitious (apparent) motion through perturbations of the B0 magnetic field caused by changes in abdominal volume when air enters the lungs (Brosch et al., 2002; Raj et al., 2001). Apparent motion of this type would be expected to be most readily visible in the phase encoding direction (Brosch et al., 2002; Durand et al., 2001; Raj et al., 2001), which is consistent with our observation that HF-motion is far more obvious in the phase-encoding direction (**Fig. 1**). Support for this mechanism has been provided by the demonstration of similar respiratory-related fluctuations in fMRI signal of phantoms placed near the head (Raj et al., 2001), indicating that they are not driven by head motion. Moreover, a mechanical model of abdominal respiration can reproduce phase-encoding modulations of the fMRI signal in a phantom at respiration rates, despite the lack of any physical connection between the two objects (Brosch et al., 2002).

The magnitude of factitious head motion is determined by sequence parameters that affect the EPI bandwidth (Hz/pixel) in the phase encoding direction (see Fair et al., 2020, for additional discussion of this point). Relevant sequence parameters include the TE and level of acceleration. In brief, shorter TEs and greater acceleration reduce sensitivity to main field perturbations (expressed as “mm/Hz”) but also reduce the EPI signal-to-noise ratio. GRAPPA acceleration may have contributed to the relatively modest level of factitious head motion in the UIUC Lifespan data (see **Fig. 2** and **Supp. Table 1**).

In a recent paper, Power and colleagues analyzed several short TR multiband datasets and suggested that there may be multiple respiration-related effects present in realignment parameters, some of which are associated with true head movements and some of which are associated with apparent “pseudo-motion” (Power et al., 2019). Spectral aliasing obscures the appearance of respiratory motion in single-band datasets (Fair et al., 2020) and precludes the detailed spectral decomposition from Power et al. (2019). However, the predominance of HF-motion in the phase-encoding direction (Fig. 1B) suggests that it is factitious (Fair et al., 2020), consistent with past work (Brosch et al., 2002; Raj et al., 2001). Regardless of their source, the motion-FC analyses that we conducted suggest that HF-motion does not bias functional connectivity in the same way as transient head movements reported in the past (Power et al., 2012; Satterthwaite et al., 2012).

It is worth noting that the current observations are distinct from other kind of respiratory effects that may need to be addressed by different methods. Apart from the factitious effects of respiration at a constant rate, changes in respiratory rate can also cause profound intermittent changes in fMRI signal (Birn, 2012; Power et al., 2017b). These effects are likely caused by alterations in arterial pCO_2_ levels that induce widespread changes in T2*-weighted signals and consequent distortions of functional connectivity measures. Global signal regression appears to be an effective strategy to address these occasional respiratory effects (Power et al., 2017b).

### Addressing HF-motion in fMRI analysis leads to data savings

While the potential contamination of fMRI by high frequency respiration-related effects has been appreciated for nearly 20 years (Brosch et al., 2002; Durand et al., 2001; Raj et al., 2001; Van de Moortele et al., 2002), the availability of multiband sequences and the typical employment of realignment parameters to denoise fMRI data has caused renewed interest in this phenomenon (Chen et al., 2019; Fair et al., 2020; Power et al., 2019) (see also popular blog posts on this topic by Jo Etzel and Ben Inglis (Etzel, 2016a, b, c; Inglis, 2016a, b)). With the shorter TRs associated with multiband data it is possible to more clearly identify respiration-related content in realignment parameters, as Nyquist limits are higher and respiration rates do not alias to lower frequencies (Chen et al., 2019; Etzel, 2016a; Fair et al., 2020; Inglis, 2016b; Power et al., 2019). Moreover, many fMRI processing pipelines use measures of frame-to-frame changes in realignment parameters as an estimate of participant head motion, and censor frames with even small amounts of head movement (e.g., 0.2 mm.) (Ciric et al., 2017; Power et al., 2012; Satterthwaite et al., 2019; Satterthwaite et al., 2012). Our motion-FC analyses (**Fig. 6**) once again demonstrated that a combination of GSR and motion censoring most effectively reduces bias in functional connectivity analyses, consistent with past reports (Ciric et al., 2017; Parkes et al., 2018; Power et al., 2012; Power et al., 2014a)^3^. However, we additionally show that high-frequency factitious motion can obscure small transient movements, disrupting the connection between motion parameters and BOLD signal disruptions. This is the case in virtually all participants in fast multiband datasets (Fair et al., 2020), but here we show that it is also relatively common in participants studied with slower single-band fMRI, especially those who are older, have higher BMI, and have lower cardiorespiratory fitness. We speculate that this increased prevalence may be due to a combination of factors that changes breathing rates, the size of breaths, and the size of the abdominal changes associated with respiration (see *previous section*).

This factitious head motion can cause substantial data loss in analyses employing FD-based frame censoring, and data loss may be exacerbated in certain populations. One question is whether this data loss is necessary, or if we can recover data contaminated by HF-motion without negatively affecting fMRI analyses. Here we show evidence in favor of the latter possibility: correcting HF-motion contamination via filtering saves substantial amounts of data without increasing motion-related biases in functional connectivity analyses. In participants without prominent HF-motion, filtering has a fairly minimal effect, with comparable frames identified for removal with and without filtering. In contrast, in participants with prominent HF-motion, large amounts of data can be recovered, and grayplots suggest that these filtered motion metrics retain a greater association with disruptions in the BOLD signal (consistent with previous results from multiband data in adolescents (Fair et al., 2020)). Furthermore, we conducted an extensive quantification of the effects of motion on functional connectivity analyses and demonstrated that filtered motion parameters still adequately correct biases, including both those that are distant-dependent (the biases most affected by motion censoring) as well as functional connectivity artifacts that are common across all regions. Further, we replicated these results in an independent dataset focused on a different population (young to middle aged neurotypical controls). Thus, our findings indicate that filtering effectively corrects motion estimates, saves large amounts of data, and adequately addresses motion confounds in functional connectivity analyses.

Motion censoring is most heavily utilized in functional connectivity pipelines but also has been increasingly adopted in task-based fMRI (e.g., see (Siegel et al., 2014)). Thus, while HF-motion is perhaps of greatest importance in FC analyses, it may also influence task-based processing strategies. The present findings suggest that filtering motion parameters would be beneficial in that context. Moreover, even processing strategies that do not explicitly censor high-motion frames may benefit from removing high frequency content from motion parameters if they do not track well with true head motion (Chen et al., 2019; Fair et al., 2020). As shown in Fig. 4, filtered motion metrics appear to retain a better correspondence with fMRI artifact, suggesting that any pipeline that includes motion parameters in nuisance regression (Ciric et al., 2017; Parkes et al., 2018; Power et al., 2015) or for identification of high quality participants (Parkes et al., 2018) will also benefit from filtering.

### Implications for studies across different populations

We show HF-motion is somewhat trait-like, with stability across sessions in a given individual. HF-motion is more prevalent in older adults, those with higher BMI, and those with lower cardiorespiratory fitness. This indicates that the current findings – and solution of using filtered motion parameters – are most relevant to experiments focused on populations with these characteristics. Thus, in datasets loaded with these characteristics (PIB-PD, SZ, Dallas, UIUC, Iowa-lesion), filtering motion parameters saved substantial amounts of data (12-19%), much more than seen in the young adults (and children) collected for the MSC and Petersen-120+ datasets. These findings indicate that effectively managing factitious head motion is particularly critical in datasets acquired in older adults, such as studies of aging, Parkinson Disease (PIB-PD), or stroke.

Interestingly, this result also provides a clue as to why HF-motion has not been highlighted in past work on motion censoring. Several of the most prominent studies done in this domain (Ciric et al., 2017; Power et al., 2011; Satterthwaite et al., 2012; Van Dijk et al., 2012) used child, adolescent, and young adult data. However, due to the younger age of these participants, it is likely that HF-motion was not as prominent (perhaps due to a combination of smaller chest sizes and faster respiration rates that alias motion into lower frequencies where it is hard to distinguish from true movements). It is only with the increasing prevalence of multiband datasets that HF-motion is clearly evident across this age range (Fair et al., 2020; Power et al., 2019).

In broad strokes, our findings concur with previous studies which have demonstrated a connection between motion metrics and age and/or BMI (Madan, 2018; Savalia et al., 2017). However, true (lower frequency) head motion may also be correlated with these demographic characteristics, even when HF-motion is explicitly removed (Siegel et al., 2017), suggesting that some participant characteristics may relate to both true and factitious motion. Older individuals especially with those with high BMI and/or lower fitness may have more trouble breathing when supine as their diaphragm is pushed down by body weight. Our results also agree with and extend upon previous findings from Power et al. (2019) demonstrating that HF-motion is heritable and related to BMI in multiband data: we replicate these results in two new datasets based on single band data and demonstrate, further, a separable relationship with age and cardiorespiratory fitness.

One question that the current findings pose is what to do for participants/datasets collected with slower TRs that exhibit less prominent HF-motion. While filtering motion parameters may not be necessary in these situations, our findings suggest that filtering motion parameters does not appreciably alter the processing of low HF-motion participants and results in a similar relationship to BOLD signal abnormalities and similar censoring masks. Thus, it may be more prudent to apply the filtering procedure across people/datasets than to selectively choose to include this processing step in only some participants. Moreover, it may be difficult to provide an exact age cut off for when to filter motion metrics, as the relationship between HF-motion and age had suggestions of a non-linear relationship (see **Supp. Fig. 4**). All three datasets examined in this manuscript showed increases in HF-motion from relatively early to mid-adulthood (ages 20-40), but also evidence of slightly decreased HF-motion in the oldest studied ages. Even in young adult datasets, we have found that unfiltered FD-based frame censoring can occasionally lead to a subject being removed when they would otherwise be saved by filtering HF-motion (e.g., participant MSC03 in the MSC dataset; Fig. 2B), which argues for including this processing step across the board. In general, these findings suggest that filtering motion parameters may be advantageous in many participants over college age and may be particularly useful for equating the fidelity of motion metrics across the lifespan.

### Other approaches to addressing HF-motion

We advocate filtering motion parameters before FD calculation to reduce unnecessary frame censoring caused by HF-motion and validate this preprocessing approach in functional connectivity analysis. However, in this section, we consider other approaches for addressing HF-motion.

One simple solution to the elevated FD baseline and high frame censoring seen with HF-motion could be to raise the FD threshold for motion censoring. However, two issues arise with this approach. First, because only a subset of individuals show prominent HF-motion, FD thresholds would either need to be changed on a subject-specific basis or risk under-censoring in participants without HF-motion. Secondly, in our experience, HF-motion raises the FD baseline, but does not particularly enlarge effects associated with typical head movements and disruptions in the BOLD signal. Thus, in the presence of HF-motion it is simply more difficult to discriminate typical head movements from the baseline, in effect making FD a “noisier” measure. Raising the FD threshold will not correct this noise issue.

Previous studies have also suggested correcting for HF-motion through filtering, but adopted a notch filter (potentially even individually selected; (Fair et al., 2020)) instead of the low-pass filter proposed here. This prior work suggests that a notch filter is an effective correction approach in multiband data. However, in slower single band data, the narrower frequency range and aliasing of respiration content means that HF signals extend over a relatively broad range of resolved frequencies, making it more difficult to select a narrow set of frequencies to remove from analysis. Selecting individual-specific ranges of frequencies to filter may also be a viable solution if good respiratory rate data is available, but it is not clear that this strategy substantially improves generic filtering in practice (Fair et al., 2020). We suggest that simple low-pass filtering is effective in slower single band fMRI.

Other approaches to correcting for head motion might also be considered in datasets with HF-motion. For example, DVARS (Smyser et al., 2010) rather than FD could be used for frame censoring (Burgess et al., 2016; Power et al., 2012). In our experience, this does at times work better than the (unfiltered) FD in the presence of HF-motion, but the effects are variable across participants (see Supplemental Material in (Burgess et al., 2016) for similar observations and potential causes for this variability). There is also some evidence that DVARS may also be affected by the same B0 signal distortions that are present in FD measures, although to a lesser degree (Fair et al., 2020). Parkes et al. (2018) also recently suggested that a combination of ICA-AROMA and global signal regression may perform approximately as well at reducing FC motion artifacts as censoring with GSR, as long as “high motion” individuals are removed from study (defined as those with < 4 min. of data after censoring; see however Ciric et al. (2017) for slightly different findings in an alternate study population). This may be a reasonable option to prevent excessive data loss in analysis. Importantly, regardless of which analysis approach is chosen (censoring through FD or DVARS, or use of ICA-AROMA with strict subject exclusion), it is imperative to have an accurate measure of subject motion to (a) correctly identify frames for censoring/spike regression, (b) correctly identify which subjects should be excluded due to high rates of motion, and/or (c) accurately remove effects of motion through nuisance regression of motion parameters, a step included in almost all analysis streams (Ciric et al., 2017; Parkes et al., 2018). Thus, our findings on HF-motion and the fidelity of fFD metrics are relevant to most current analysis pipelines, regardless of the precise censoring criteria adopted.

Another exciting avenue for future research is the possibility of dynamically correcting for homogeneity distortions caused by respiration. A procedure of this type has been described for diffusion tensor imaging (Andersson et al., 2018), which also is based on EPI, but no such procedure has yet been developed for fMRI, as far as we are aware. In contrast, current homogeneity field map correction approaches for fMRI are static (i.e., a single correction is applied to a full functional run), and are typically applied after motion realignment. Thus, these current approaches will not correct for the dynamic frame-to-frame distortions caused by respiration.

### Strengths and Limitations

We were able to demonstrate the characteristics of HF-motion, likely due to respiration, and their association with different participant factors across multiple diverse populations, scanner sequences, and sites. Moreover, we proposed an approach to address HF-motion that could lead to significant additional data retention (again, replicated across multiple datasets). Finally, we conducted a rigorous analysis of the consequences of HF-motion on FC analysis, demonstrating that our filtering approach did not reintroduce motion contamination into FC.

However, none of the datasets analyzed in the current study included respiratory measures. Therefore, we cannot conclusively demonstrate that HF-motion in these datasets is respiratory-related. Instead we rely on inference from other related studies (Fair et al., 2020; Power et al., 2019).

A second limitation is that some of the findings were not consistent across datasets. For example, the typical frequency of HF-motion varied across datasets. These differences may have been associated with different baseline respiration rates and body sizes across participants (who were of different ages and fitness levels). Given our lack of respiratory data it is not possible to exclude other causes. In addition, differences in TR (from 2-2.5) likely would alter the aliasing of the respiratory signal to slightly different frequencies across datasets. The relationship between HF-motion and age was also only present in two out of three datasets tested with a linear approach. However, encouragingly, the two consistent datasets showed a very similar magnitude of relationship and all three datasets showed upward trends with age when using non-linear fits (**Supp. Fig. 4**) along with slight downward trends in the oldest ages. We speculate that as studies recruit participants into older ages, they may increasingly select for a particularly healthy/active subset of older adults, and this may have been exacerbated in the UIUC Lifespan dataset due to its smaller size and multi-session nature. Additional findings related to sex and diagnosis (and interactions between all of these different factors; reported in the **Supp. Table 3** and **Supp. Fig. 5**) were also inconsistent across datasets. Future studies will be needed to better understand these aspects of variation.

Finally, given the retrospective nature of this work, we were only able to examine relationships between HF-motion and demographic measures that were collected in each dataset. In all cases this included age and sex, but only in two datasets (Dallas Lifespan, UIUC Lifespan) was BMI data collected, and only one dataset included additional measures of cardiorespiratory fitness (UIUC Lifespan). Moreover, age, BMI, and cardiorespiratory fitness only explain a moderate amount of the variance in HF-motion. Thus, in addition to these variables, it is likely that other demographic/behavioral factors (such as baseline respiratory rates, levels of anxiety in the scanner - which may, in turn, affect respiratory rates – and lung volume) also influence HF-motion. It will be interesting to examine HF-motion in additional studies/datasets that target collection of these additional measures.

## Conclusions

We analyzed multiple datasets to demonstrate that fMRI motion measures can be contaminated by factitious respiratory effects, even in conventional single-band fMRI. This contamination was strongest in the phase encoding direction and was relatively stable over sessions, which, together with past reports, is consistent with a relationship to respiration. Moreover, the high frequency contamination varied with subject characteristics, being more prominent in older adults, those with higher BMI, and those with lower cardiorespiratory fitness. We demonstrate that filtering motion parameters corrects for this high frequency modulation and saves substantial amounts of data in most datasets, while still adequately addressing motion biases in functional connectivity estimates. These results suggest incorporating HF-motion filtering will improve analysis of fMRI datasets, especially those acquired from older and less fit individuals.

## Acknowledgements

The authors thank Dr. Hongyu An for helpful discussion of fMRI sequence parameters. Funding was provided by NIH grants MH118370 (CG), F32NS092290 (CG), NS075321 (JSP), NS097437 (MCC), NS058714 (JSP), NS098577-01 (AZS), MH066031 (DMB), AG059878 (MF, GG), R56MH097973 (GG; MF), R25MH112473 (TOL), MH096773 (DAF), MH091238 (DAF), MH115357(DAF), DA041148 (DAF), MH122066 (DAF; NUFD), NS088590 (NUFD), TR000448 (NUFD), AG063930 (GSW), as well as a McDonnell Foundation Collaborative Activity Award (SEP, DT), a James S. McDonnel Foundation Understanding Human Cognition Award (GSW), pilot funding support from the Northwestern Alzheimer’s Disease Center (NIA P30AG13854 to CG), an award from the Gates Foundation (DAF), the Destafano Innovation Fund (DAF), the Jacobs Foundation (NUFD), an OHSU Fellowship for Diversity and Inclusion in Research Program (OM-D), a Tartar Trust Award (OM-D), the OHSU Parkinson Center Pilot Grant Program (OM-D) and awards from the American Parkinson Disease Association (APDA) Advanced Research Center for PD at WUSTL; Greater St. Louis Chapter of the APDA; McDonnell Center for Systems Neuroscience; Washington University Institute of Clinical and Translational Sciences (MCC), Barnes Jewish Hospital Foundation (Elliot Stein Family Fund), and the Riney Foundation (JSP). The authors would like to thank Denise Park for providing access to the Dallas Lifespan Brain Study data, collected under NIH grant 5R37AG-006265-25.

## Footnotes

1 The Nyquist folding frequency is one half the sampling rate.

2 Note that for simplicity, the censoring mask computations for **Fig. 4 and 5, Table 2**, and related sections of the text are made with a simple threshold mask (e.g., FD<0.2 or fFD<0.1), but not including other elements of censoring that we typically add to FC processing (Power et al., 2014a). These other elements include: censor frames at the start of each run (14 frames) and frame segments that are less than 5 (or 3 – see **Supp. Fig. 13**) contiguous units long.

3 As a cautionary note, our motion-FC analyses also demonstrated that bias reduction is sensitive to the minimum number of frames present across participants (**Supp. Fig. 13**), consistent with reports from Parkes et al. (2018). A minimum of 150 frames (∼5.5 min.) was necessary to strongly alleviate distance-dependent motion bias in FC. In contrast, reducing the minimum frame number to 100 or 50 progressively increased the distance-dependent bias present in the data. Thus, these analyses suggest ensuring a relatively higher (5 min.) frame minimum for maximizing bias reduction in functional connectivity analysis.

