## Supplemental for "Removal of high frequency contamination from motion estimates in single-band fMRI saves data without biasing functional connectivity"

### SUPPLEMENTAL MATERIALS

#### Supplemental Methods

Iowa-Lesion Dataset, Participant Characteristics and MRI parameters:
The Iowa-Lesion dataset consisted of 28 participants (15 F, ages 25 – 78) with focal brain lesions collected at the University of Iowa. Participants were identified from the Iowa Lesion database and partially overlap with the participants reported in Warren et al. (2014). Each participant completed 1-3 sessions of MRI data collection on a Siemens 3T Trio.

MRI data collection included T1 MPRAGE, T2, field maps, and resting-state functional EPI scans (eyes open fixation). As only data associated with the functional EPI scans was analyzed here, we focus on those parameters: TR = 2.26 s., TE = 30 ms., flip angle = 80 degrees, 36 slices collected at 4mm. slice thickness with a matrix size = 64 x 64 (voxel size = 3.44 x 3.44 x 4 mm.), with interleaved slice acquisition. Each resting state run was 216 TRs long and participants completed 2-10 resting state runs.

PIB-PD Participant Selection Criteria:
The PIB-PD dataset is drawn from a large longitudinal dataset collecting MRI, PET, CSF, and behavioral measures from Parkinson participants and age-matched controls (see Gratton et al. (2019) for original participant inclusion criteria). At each longitudinal fMRI session, participants completed 1-3 resting-state fMRI runs of 200 frames each, depending on participant comfort levels. Only the first session of each participant with sufficient data for functional connectivity analyses (see criteria below) was used here.

In particular, in this dataset, we began with the 153 participants from Gratton et al. (2019) and added in 37 additional participants (originally excluded due to movement, dementia/diagnostic criteria, or who had not yet been collected and processed). Of this set of 190 individuals, we included any participants who passed a relaxed criteria of at least 30 frames in a single run after censoring using all three of the different censoring criteria employed in this manuscript (see *Methods: FD vs. fFD censoring*). This resulted in a set of 143 final participants. All of these participants were used for motion parameter calculations and characterization. Some subjects were further removed from functional connectivity analyses based on specific criteria described in that section, in order to match participants across FD-QC analyses.

Cardiorespiratory fitness
Cardiorespiratory fitness is most accurately measured using VO2max during exercise. However, older or lower-fit individuals may have conditions that prevent them from safely participating in these stressful exercise tests. Thus, CRF was estimated using an equation shown to be highly predictive of VO2max not only in large samples above 10,000 participants (Jurca et al., 2005; Stamatakis et al., 2013) but also in small samples (McAuley et al., 2011) and when only older adults ranging from ages 60-80 were used (Mailey et al., 2010). Specifically, CRF was estimated by a linear combination of weighted variables using the equation: CRF = Gender (2.77) −Age (0.10) −BodyMass Index (0.17) − Resting Heart Rate (0.03) + Self-Reported Activity Score + 18.07 (Jurca et al., 2005).

Functional Connectivity (FC) Analysis of the PIB-PD dataset
*Structural image processing and alignment:* Functional data was aligned to a T1 structural image from each participant using affine registration. The T1 image was also aligned to a Talairach group template using a linear affine registration procedure. These transforms were concatenated and applied to the functional data in a single step (along with resampling to 3mm. isotropic resolution). Freesurfer segmentation of the structural T1 data was used to create white matter and cerebrospinal fluid nuisance masks for each individual.

*FC Preprocessing*: The following steps were conducted on the functional data, following Power et al. (2014) to remove sources of noise. (1) Data was demeaned and detrended. (2) Nuisance regression was conducted, regressing out BOLD activity related to global signal, white matter, cerebral spinal fluid, the six motion parameters, their derivatives, and the first and second order Volterra expansions of the motion parameters. (3) High motion frames (see Methods "Identification of high motion frames" for criteria) were removed and interpolated over using a spectral interpolation procedure that preserved the frequency content of the underlying data. (4) Bandpass temporal filtering (0.009 – 0.08 Hz.) was applied to the data. (5) Spatial smoothing (6mm. FWHM) was applied to the data. After processing, interpolated frames were removed ("censored") from the timeseries. Additional analyses were conducted in which either censoring, global signal regression, or both were eliminated from the processing stream to examine how these steps affected FC.

*Grayplots:* A modified version of the Power 2018 grayplots was generated to examine the relationship between FD, fFD and gray matter signal changes. These plots are shown after step 2 above (that is, after demeaning and detrending as well as nuisance regression). Note that while grayplots are useful tools for visualization, the visibility of deflections of different types may be affected by the ordering of BOLD timeseries in grayplots (Aquino et al., 2019). In this manuscript, grayplot ordering is semi-random as is common in previous grayplot depictions (Power, 2017).

*FC Calculation:* Functional connectivity was calculated among a set of 300 spherical regions spanning cortex, subcortex, and the cerebellum ((Seitzman et al., 2019); these regions represent an expansion of the 264 regions from Power et al. (2011) with better subcortical coverage). Denoised and censored timeseries were averaged within each region and then correlated between pairs of regions using Pearson correlation.

### Supplemental Tables

| Dataset | Site | Scanner | TR (s) | TE (ms) | Flip angle (deg) | Slice number | Voxel size (mm) | Acceleration (GRAPPA, iPAT, partial Fourier) | Run length (TRs) | Number of runs |
|---|---|---|---|---|---|---|---|---|---|---|
| **PIB-PD*** | WUSTL | Siemens Trio 3T | 2.2 | 27 | 90 | 48 | 4.0 x 4.0 x 4.0 | n/a | 200 | 1-3 |
| **Barch-SZ** | WUSTL | Siemens Trio 3T | 2 | 27 | 90 | 35 | 4.0 x 4.0 x 4.0 | n/a | 221 | 8 (task) |
| **Petersen 120+** | WUSTL | Siemens Trio 3T | 2-2.5 | 27 | 90 | 32-36 | 4.0 x 4.0 x 4.0 | n/a | 76-724 (avg: 170) | 1-6 (avg: 2.4) |
| **MSC** | WUSTL | Siemens Trio 3T | 2.2 | 27 | 90 | 36 | 4.0 x 4.0 x 4.0 | n/a | 818 | 10 |
| **Iowa Lesion** | U Iowa | Siemens Trio 3T | 2.26 | 30 | 80 | 36 | 3.44 x 3.44 x 4.0 | n/a | 216 | 2-10 |
| **Dallas Lifespan** | UT Dallas | Phillips Achieva 3T | 2 | 25 | 80 | 43 | 3.4 x 3.4 x 3.5 | n/a | 154 | 1 |
| **UIUC Lifespan** | UIUC | Siemens Trio 3T | 2 | 25 | 90 | 38 | 2.6 x 2.6 x 3.0 | GRAPPA factor 2 | 180 | 2 |

**Supplemental Table 1:** Summary of functional MRI parameters from the seven datasets analyzed in this manuscript. All datasets were collected with gradient echo EPI sequences and A/P phase encoding. Avg = average. *Primary dataset

| Dataset | Analysis Package | Alignment algorithm | Alignment reference |
|---|---|---|---|
| **PIB-PD*** | 4dfp | cross-realign3d | middle frame |
| **Barch-SZ** | 4dfp | cross-realign3d | middle frame |
| **Petersen 120+** | 4dfp | cross-realign3d | middle frame |
| **MSC** | 4dfp | cross-realign3d | Middle frame |
| **Iowa Lesion** | 4dfp | cross-realign3d | middle frame |
| **Dallas Lifespan** | SPM8 | spm_realign | first frame |
| **UIUC Lifespan** | SPM12 | spm_realign | first frame |

**Supplemental Table 2:** Summary of realignment approaches used to analyze the seven datasets in this manuscript. Default rigid body realignment options were used in each case. . Links to the relevant software packages (scripts and their documentation): 4dfp: https://sites.wustl.edu/nillabs/4dfp-documentation/; SPM8: https://www.fil.ion.ucl.ac.uk/spm/software/spm8/; SPM12: https://www.fil.ion.ucl.ac.uk/spm/software/spm12/. All packages optimize the same objective function (the spatial correlation between each frame and the reference frame).

| | WUSTL – all | Petersen 120+ | Barch-SZ | PIB-PD | Dallas Lifespan | UIUC Lifespan |
|---|---|---|---|---|---|---|
| **N** | 667 (407 F)<br>Ages 5 - 90 | 587 (359 F)<br>Ages 5 - 40 | 97 (38 F)<br>HC = 40 (20 F)<br>SZ = 57 (18 F)<br>Ages 20 - 55 | 143 (72 F)<br>HC = 41 (24 F)<br>PD = 102 (48 F)<br>Ages 46-90 | 403 (258 F)<br>Ages 20 - 89 | 49 (25 F)<br>Ages 18-75 |
| **dataset** | **0.0000** | - | - | - | - | - |
| **diagnosis** | - | - | **0.0454** | **0.0231** | - | - |
| **sex** | 0.101 | **0.0026** | 0.5813 | 0.9684 | 0.8509 | 0.0610 |
| **age** | **0.0079** | **0.0005** | 0.2596 | 0.9753 | **0.0000** | 0.5218 |
| **BMI** | - | - | - | - | **0.0000** | 0.0747 |
| **interactions** | age*sex: 0.9255<br>**age*dataset: 0.0172**<br>**dataset*sex: 0.0044** | age*sex: 0.2384 | age*sex: 0.0137<br>diagnosis*age: 0.0284<br>**diagnosis*sex: 0.0013** | age*sex: 0.8722<br>diagnosis*age: 0.4052<br>diagnosis*sex: 0.8979 | age*sex: 0.1667<br>bmi*age: 0.1971<br>bmi*sex: 0.1140 | age*sex: 0.370<br>bmi*age: 0.3935<br>bmi*sex: 0.3798 |

**Supplemental Table 3:** P-values from ANOVAs relating participant characteristics to subject motion. Each column (different dataset) represents a separate model, with rows showing participant factors included in the regression (- indicates factor was not included). All factors were z-scored before entry into the model. P-values surviving FDR correction for the number of tests (columns) are bolded (p(FDR)<0.05). *Note that the WUSTL-all regression only includes data from healthy control participants, as diagnosis status was tested separated in the Barch-SZ and PIB-PD datasets. Note also that not all participants in the Dallas Lifespan study had BMI measures; the model was conducted on the subset of participants with these measures.*

| | *FD<0.2* | *fFD<0.1* | *fFD<0.08* |
|---|---|---|---|
| ***Mean*** | 377.8 | 460.8 | 403.3 |
| ***Min*** | 152 | 241 | 190 |
| ***Max*** | 539 | 557 | 550 |

**Supplemental Table 4:** The mean, min, and max frames retained for the different censoring strategies associated with the QC-FC analyses reported in **Figure 6**. Note that these analyses were restricted to a set of N = 79 PIB-PD participants (52 PD and 27 HC) who had > 150 frames under all three frame censoring criteria (FD<0.2, fFD<0.1, fFD<0.08) with censoring masks which included removal of 14 frames at the start of each scan as well as censoring of small segments less than 5 frames long (Power et al., 2014).

**Supplemental Figures**

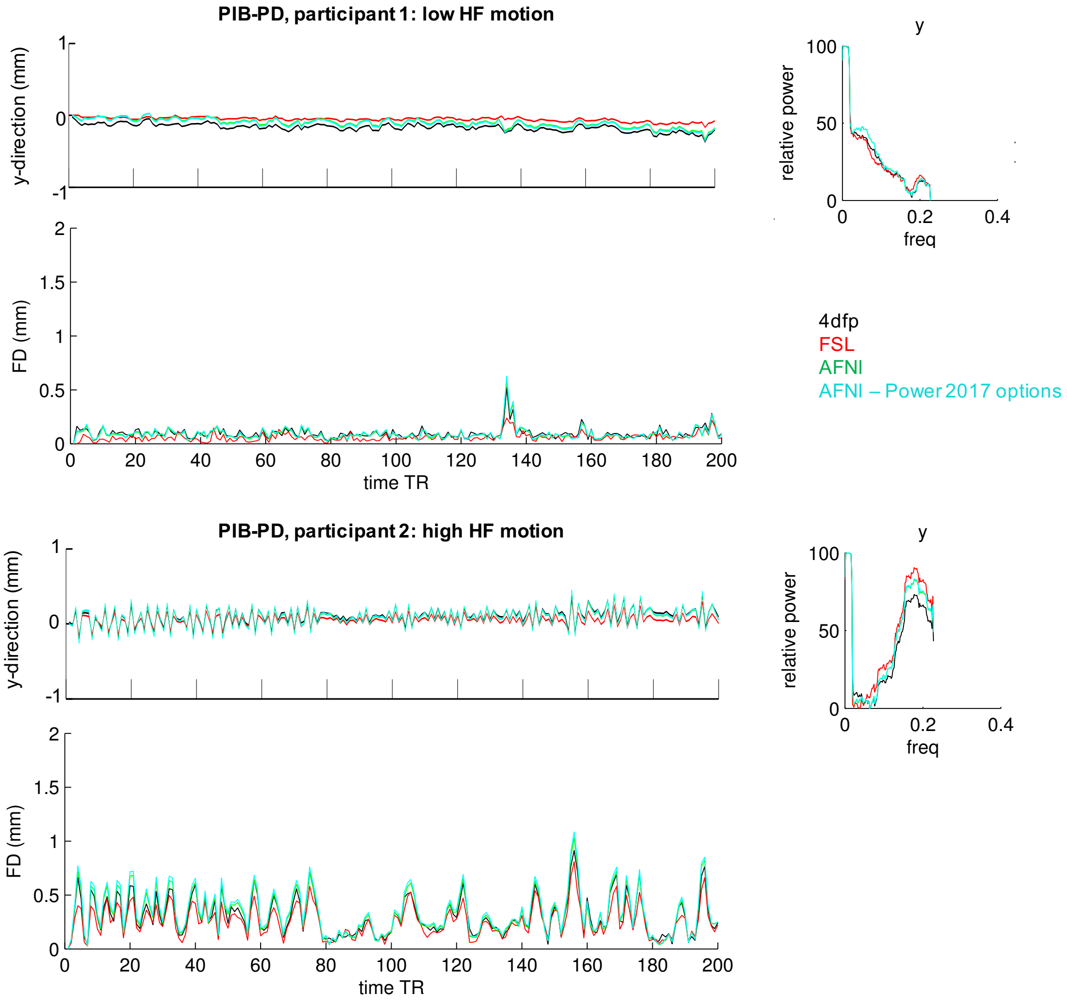


**Supplemental Figure 1:** Comparison between motion parameters from alignment algorithms from different imaging packages. Two subjects from Figure 1A are plotted again here after analysis with 4dfp (as in the manuscript), FSL's mcflirt (version 5.0) and AFNI's 3dvolreg (version 17.2.09). In all cases default parameters were used with the exception of forcing algorithms to align to the first frame to facilitate visual comparison across packages. In addition, a fourth comparison was run with AFNI using the following 3dvolreg parameters as specified in Power et al. (2017): maximum iterations 25 (-maxite 25) and cubic interpolation (-cubic). For each subject, the upper left plot shows the y-direction (phase encoding) motion traces, the upper right plot shows the relative power in these traces, and the lower left plot shows the FD parameters. There is substantial agreement across packages. Note in particular that all three packages show a lack of HF motion in subject 1 and the presence of HF motion in subject 2. Reasons for inter-package discrepancies differences contributing to the evaluation of the objective function in question (i.e. pre-blurring, field-of-view masking, etc.) and differences in algebraic conventions. Specifically, what is labeled displacement vs. rotation depends on the location of the coordinate origin (see Appendix A in (Fair et al., 2020)).

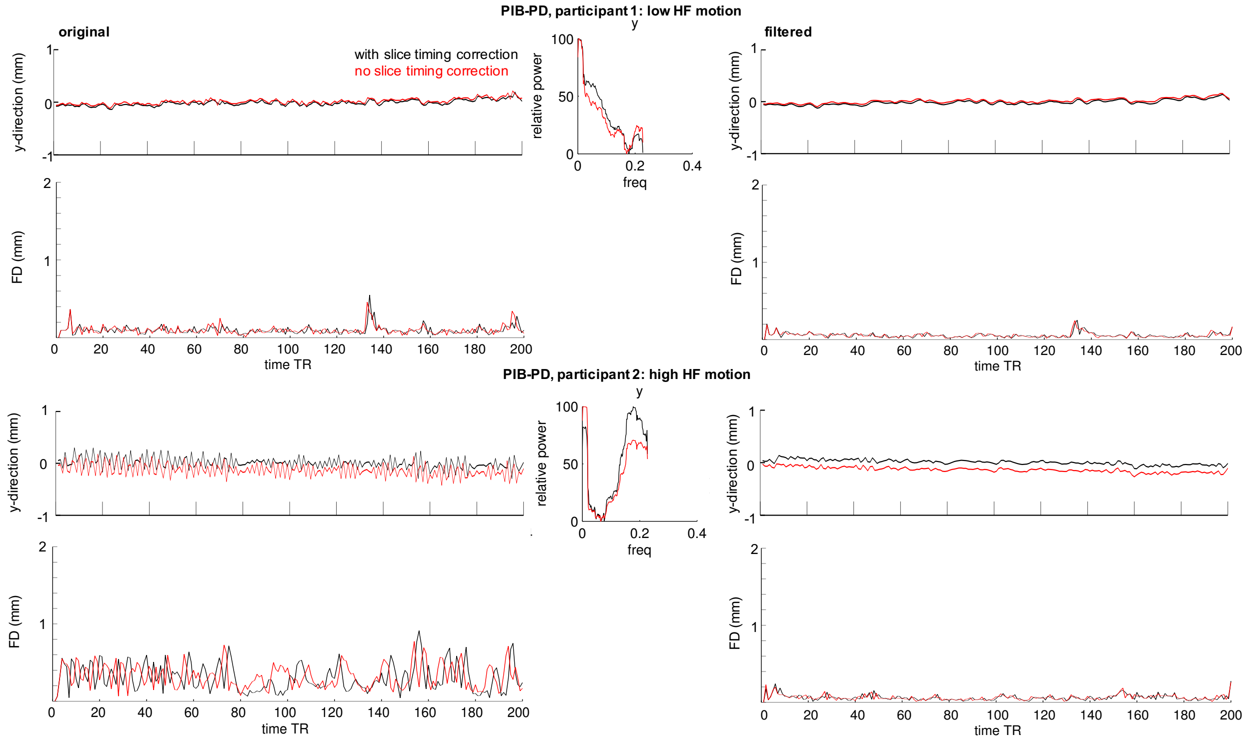


**Supplemental Figure 2:** Motion traces and FD calculated before and after slice-timing correction (STC). The top half of the panel shows data from the low HF-motion participant from Figure 1, the bottom half of the panel shows data from the high HF-motion participant from Figure 1. The left side of the figure shows the original y-direction and FD trace; the middle panel shows the relative power spectrum of the y-direction trace. The right side of the figures shows the y-direction and FD traces after low-pass filtering at 0.1 Hz. HF-motion can be seen regardless of STC. However, STC has a substantial effect on the y-direction and FD trace pre-filtering. Additional subjects and the relationships between FD with and without STC are shown in ***Supplemental Figure 3***.

Summarizing the results across these two figures, we find that (1) The FD trace is sensitive to whether or not STC correction is done. This is true for total FD as well as the y-translation component of FD. It should be noted that when STC is included, the BOLD data will also be translated in time, so differences in censored frames may be appropriate. (2) When almost all "head motion" is factitious (i.e., due to respiration), the phase-encoding direction and FD are very sensitive to STC. This is understandable as chest wall motion is near the Nyquist limit for single band fMRI. This means that small slice timing differences generate very large phase differences and substantial differences in respiratory motion amplitude. (3) Addressing HF-motion through filtering reduces the discrepancy between STC and non-STC motion traces. (4) Filtered motion parameters are effective at reducing motion biases in functional connectivity (even when computed post-STC; Figure 6).

A

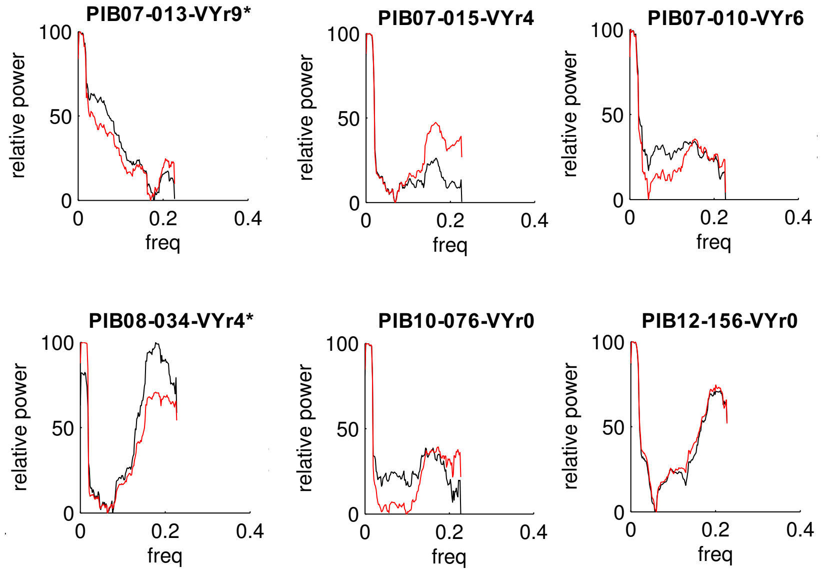


*used in previous figures as examples*

B

| | FD correlation (STC, noSTC) | fFD correlation (STC, noSTC) |
|---|---|---|
| PIB07-013-VYr9* | 0.579 | 0.830 |
| PIB07-015-VYr4 | 0.506 | 0.873 |
| PIB07-010-VYr6 | 0.567 | 0.920 |
| PIB08-034-VYr4* | -0.042 | 0.695 |
| PIB10-076-VYr0 | 0.312 | 0.761 |
| PIB12-156-VYr0 | -0.137 | 0.916 |

**Suplemental Figure 3:** Comparison of motion and FD traces before and after slice-timing correction (STC) in a set of six subjects. (A) Power spectra of the y-direction motion trace before and after STC in six subjects. Power spectra differ somewhat with STC, but HF motion can be seen in a subset of subjects regardless of STC. (B) The correlation between FD traces before and after STC, for the original FD trace (left column) or the filtered FD trace (right column). Filtering increases the relationship between FD traces.

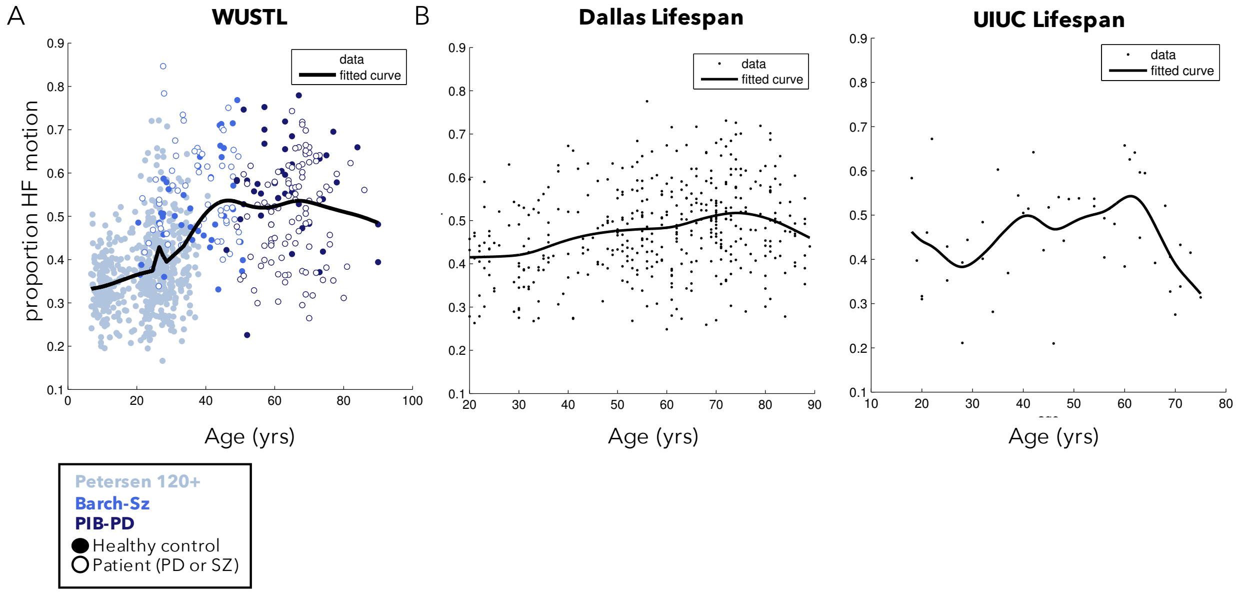


**Supplemental Figure 4:** Relationship between age and relative HF-motion in the phase encoding parameter for the (A) combined WUSTL groups (Petersen 120+, Barch-SZ, and PIB-PD), (B) Dallas Lifespan dataset, and (C) UIUC Lifespan dataset as in **Figure 3** of the main text. A smoothed spline fit is included on top of the scatterplots (smoothing parameter set to 0.0005 for the WUSTL and Dallas datasets and to 0.02 for the UIUC dataset to compensate for its smaller size).

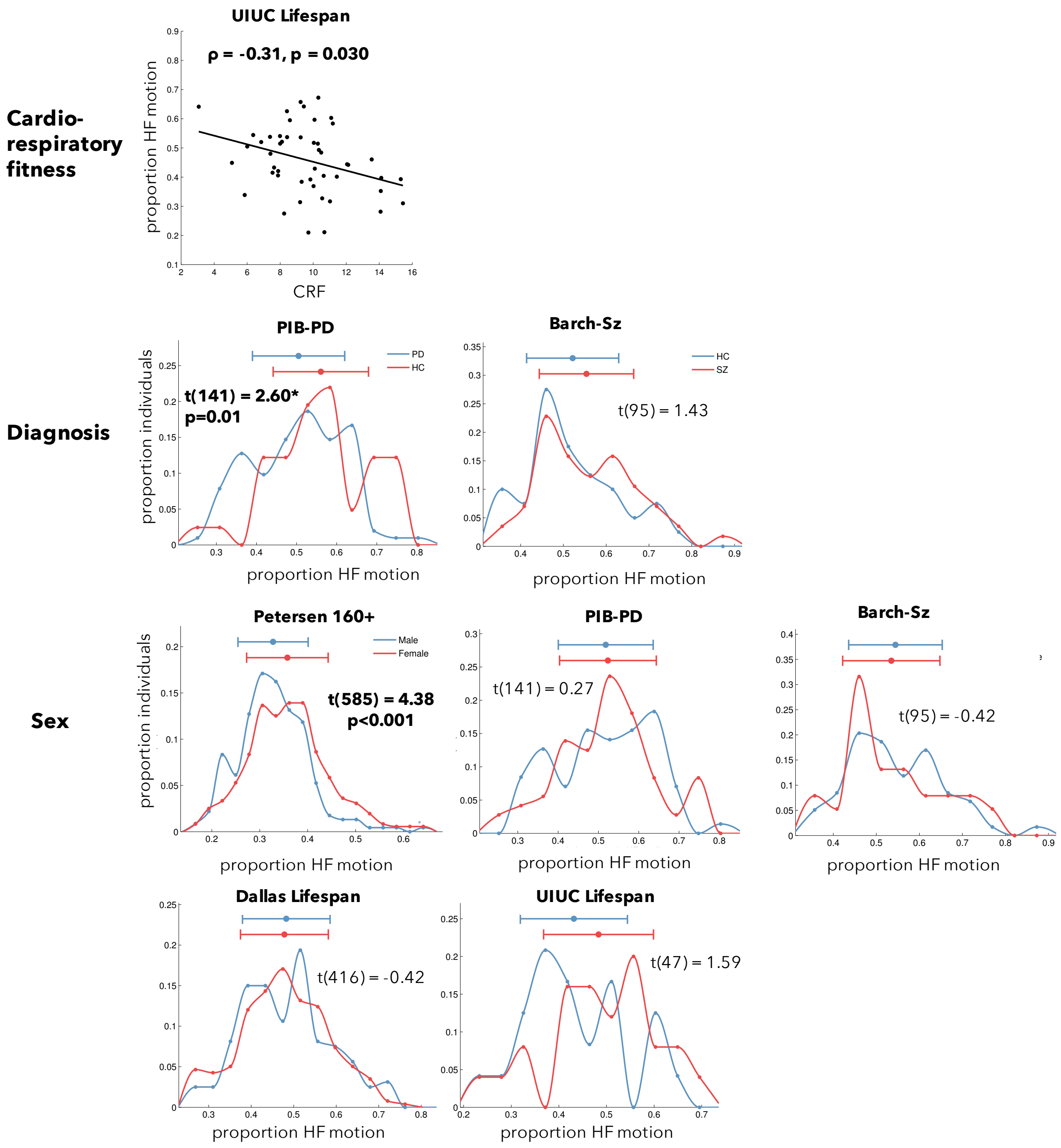


**Supplemental Figure 5:** This figure displays how the proportion of HF-motion in the phase encoding direction is related to additional participant characteristics such as cardiorespiratory fitness (top row), diagnosis (middle), and sex (bottom rows). Cardiorespiratory fitness is shown as a scatter plot and significance was tested with Spearman correlation. Sex and diagnosis are plotted as histograms and significance was tested with two-tailed t-tests. Bolded tests were significant after FDR correction for the number of tests (p(FDR)<0.05).

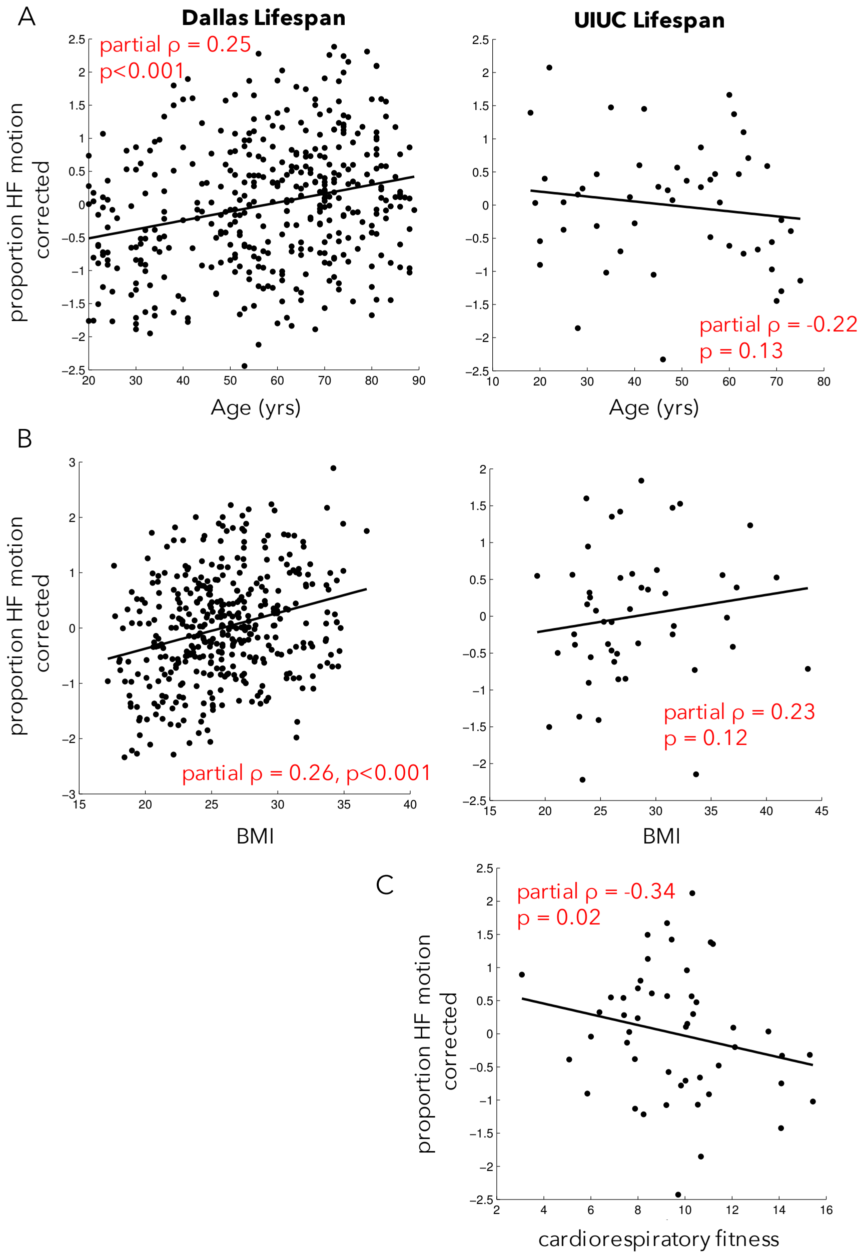


**Supplemental Figure 6:** Scatter plots of the relationship between participant characteristics and HF-motion after correcting for alternate demographic variables (e.g., age/BMI for the Dallas Lifespan dataset and age/BMI/cardiorespiratory fitness for the UIUC Lifespan dataset). (A) Relationships between HF-motion and age, (B) Relationships between HF-motion and BMI, (C) Relationships between HF-motion and cardiorespiratory fitness.

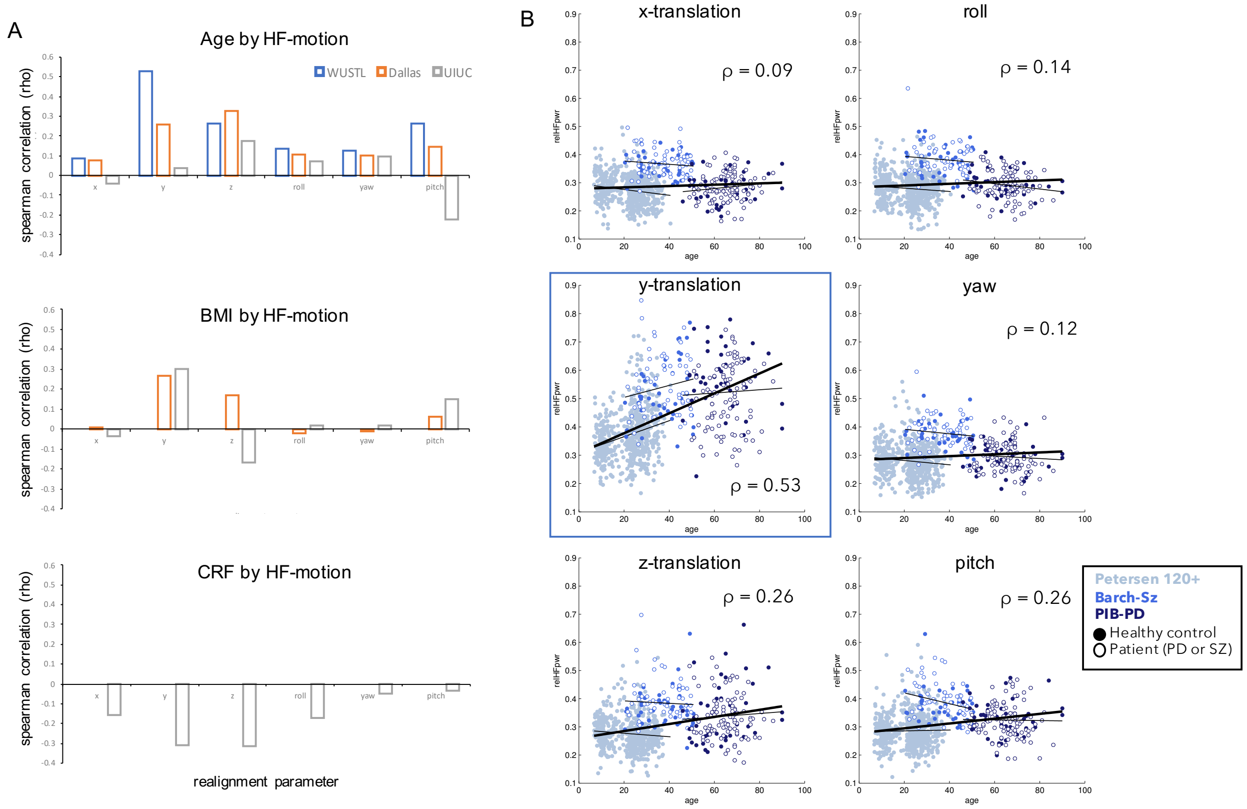


**Supplemental Figure 7:** Relationship between demographic characteristics and HF-motion across all six realignment parameters. (A) This panel shows the Spearman correlation between HF-motion in each realignment parameter and age (top), BMI (middle), and cardiorespiratory fitness (CRF, bottom) for each dataset that contains these variables. Note that the datasets differ substantially in size (WUSTL-all N = 667, Dallas Lifespan N = 403, UIUC Lifespan N = 49). (B) Scatterplots of the relationships between age and HF-motion for the WUSTL combined datasets. As can be seen, the y-translation direction (from main text Figure 3A) shows the widest range of HF-motion values and the highest correlation with age.

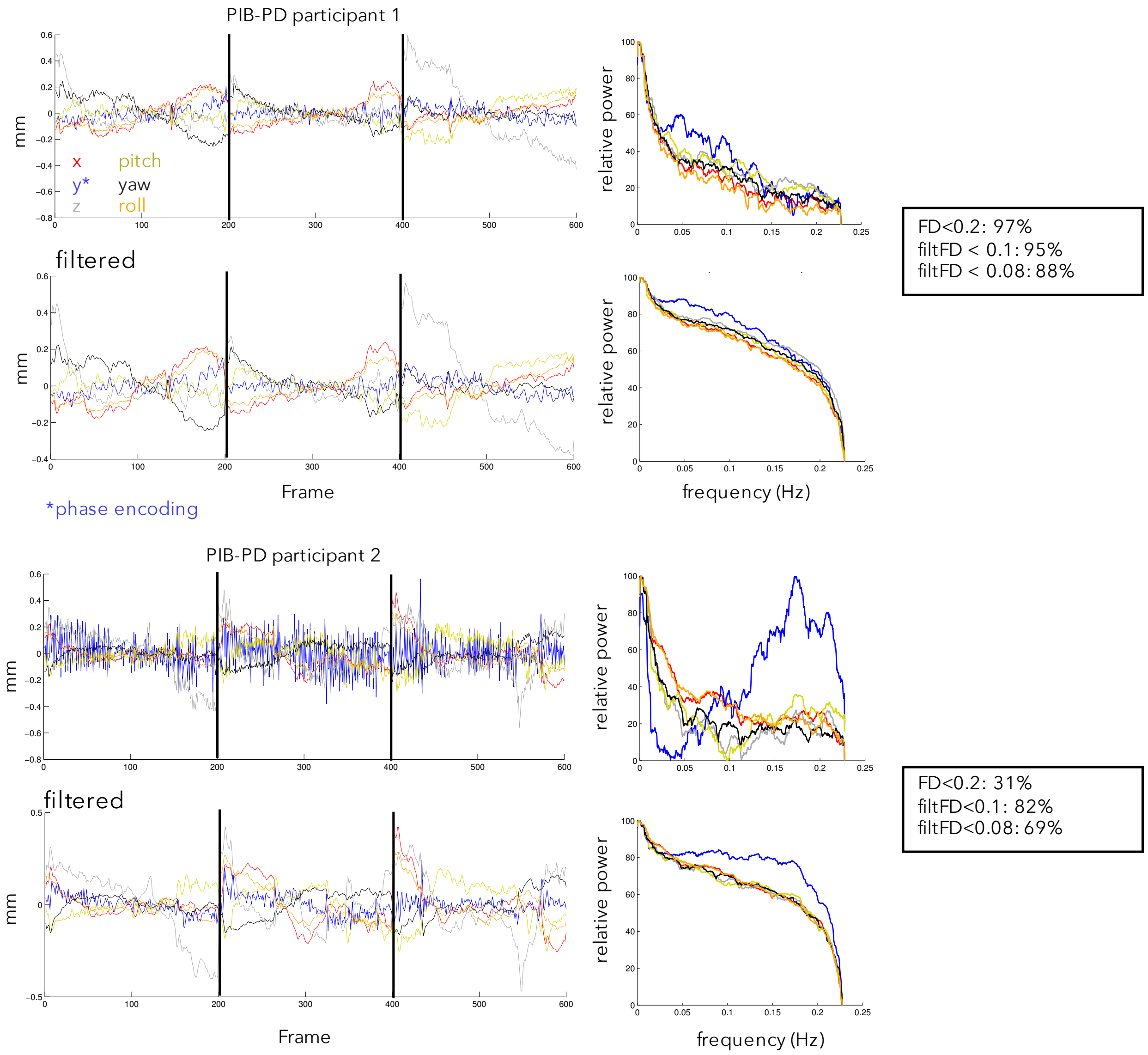


**Supplemental Figure 8:** Filtered and unfiltered motion parameters for the same participants shown in Fig.1. Left hand plots show the original motion traces over time, right hand plot show the power spectra of the traces. Boxes report the number of frames that pass the three censoring criteria tested in this manuscript.

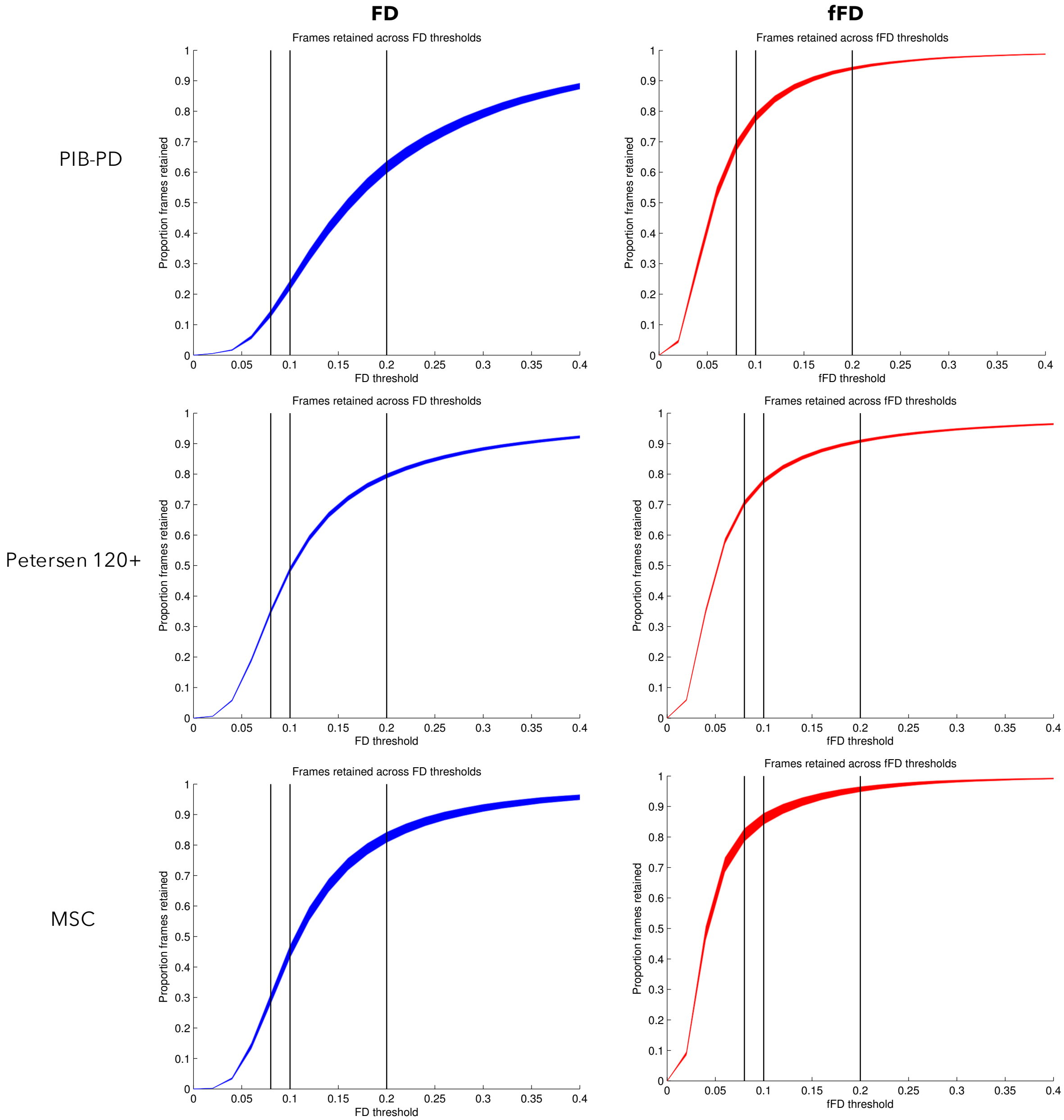


**Supplemental Figure 9:** Selecting fFD thresholds. We adopted a multi-pronged approach for selecting a threshold for the fFD measure. (1) In low motion subjects, we examined the apparent baseline of the fFD trace. The baseline of the fFD trace appeared slightly reduced relative to the baseline of the FD trace, in the absence of transient or HF-motion (e.g., see **Fig. 4A**). This examination suggested that an fFD threshold of 0.1mm would capture transient spikes in the fFD trace. (2) We examined the proportion of frames retained for subjects across a range of thresholds, using either the standard FD measure (left column above, blue) or the filtered FD measure (right column, red; shaded areas = +/- 1 standard error of the mean). In datasets with relatively little HF-motion (Petersen 120+, MSC; bottom two rows), an FD threshold of 0.2 (right-most vertical black line, left column) approximately intersects with an inflection point in frames retained, leading to ~80% of frames retained on average. In these datasets, a similar level of frames retained is achieved with an fFD threshold of 0.08 or 0.10 (first two vertical black lines, see right column). Similar levels are seen in the PIB-PD dataset (top row) with the fFD threshold (right), though note the dramatically reduced frames retained across all thresholds with the standard FD measure (left). A similar pattern is seen in the four other datasets in this manuscript (**Supp. Fig. 10**). Note also that the tmasks produced by setting fFD<0.1 and FD<0.2 are very similar to one another, especially in low motion subjects (**Fig. 5C**). (3) In subsequent analyses, we more formally benchmarked the performance of these fFD censoring thresholds at removing bias in functional connectivity (**Fig. 7, Supp. Fig. 13-15**); again, these thresholds performed as well or better than the standard FD<0.2 criteria.

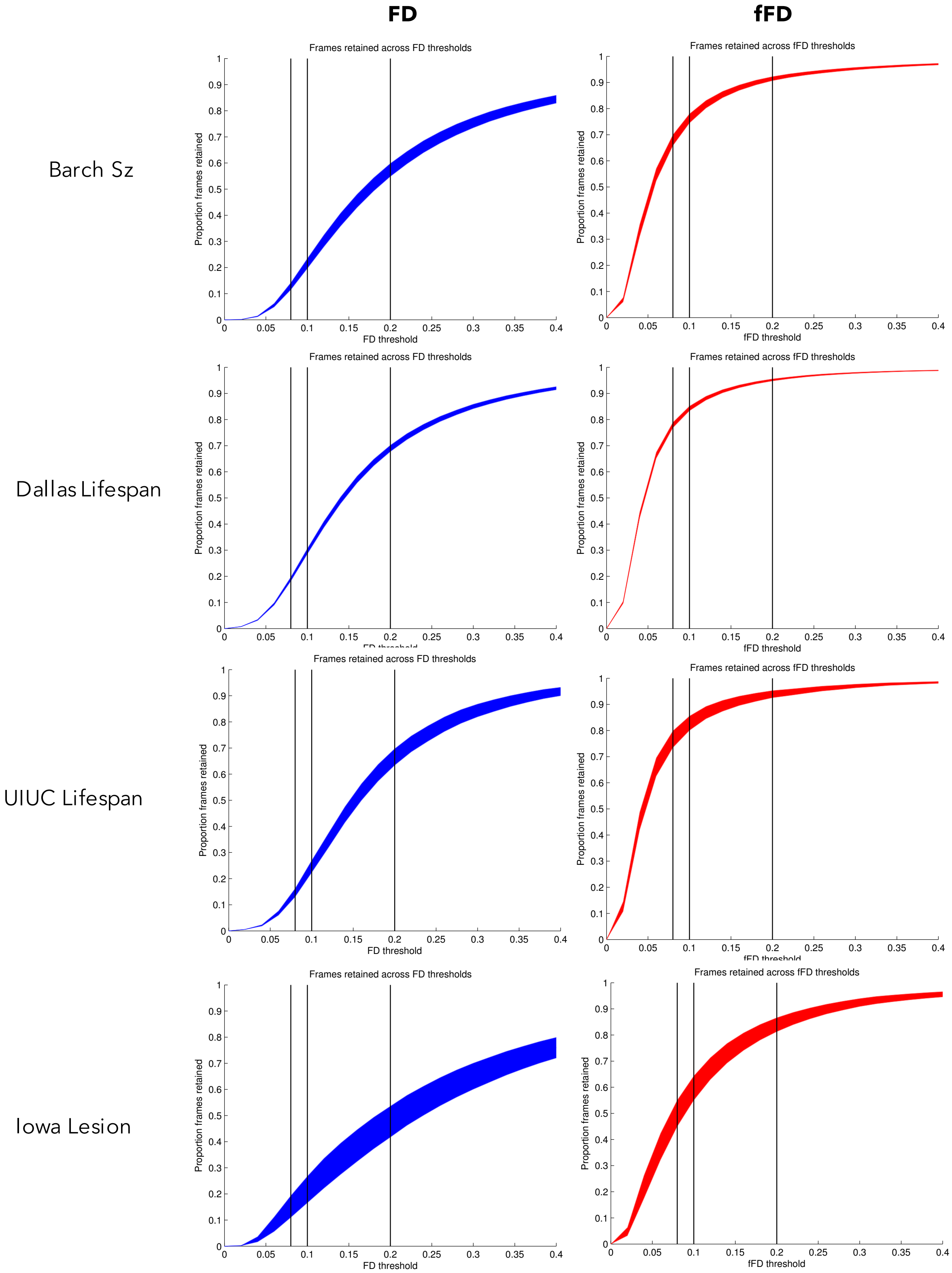


**Supplemental Figure 10:** Frames retained across FD (left column, blue) and fFD (right column, red) censoring thresholds for the four other datasets in this manuscript (rows). Note that a fFD threshold of 0.08 or 0.1 appears to intersect with an inflection point in frames retained in most datasets. Note also that the standard FD threshold of 0.2 leads to substantial data loss in all of these datasets, whose participants exhibited substantial HF-motion.

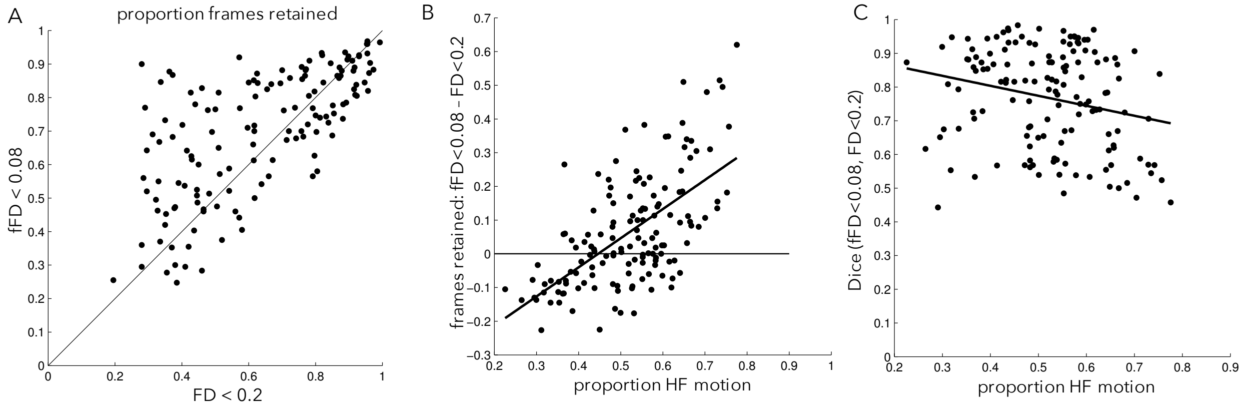


**Supplemental Figure 11:** The same comparisons made in Figure 5, but with the more conservative fFD<0.08 criteria. (A) The relationship between the number of frames retained with standard (FD<0.2) vs. this conservative (fFD<0.08) criteria. Even with this more conservative criteria, many participants show substantial data savings when motion parameters are filtered before FD calculation. (B) The proportion of HF motion is related to the gain in frames retained after censoring with fFD<0.08 criteria. (C) Censoring masks are similar between the filtered and unfiltered FD censoring criteria, especially for participants with low amounts of HF motion.

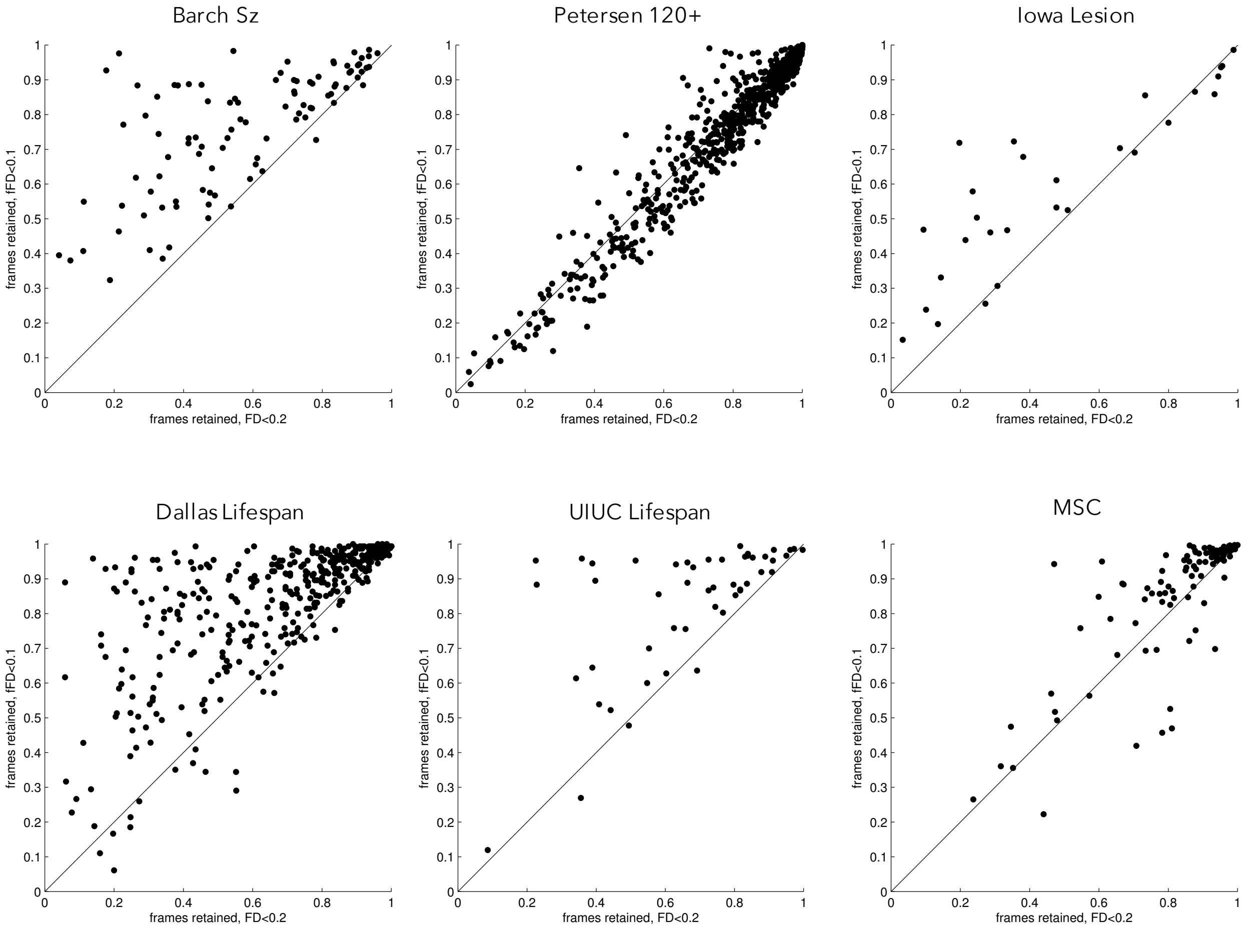


**Supplemental Figure 12:** Frames retained with standard (FD<0.2, x-axis) censoring criteria vs. censoring criteria that corrects for HF motion (fFD<0.1, y-axis) across the other six datasets examined in this manuscript. Note that despite the lower threshold, censoring saves data from censoring in a proportion of participants in each dataset, although data savings are smaller in the datasets focused on younger participants (Petersen 120+, MSC).

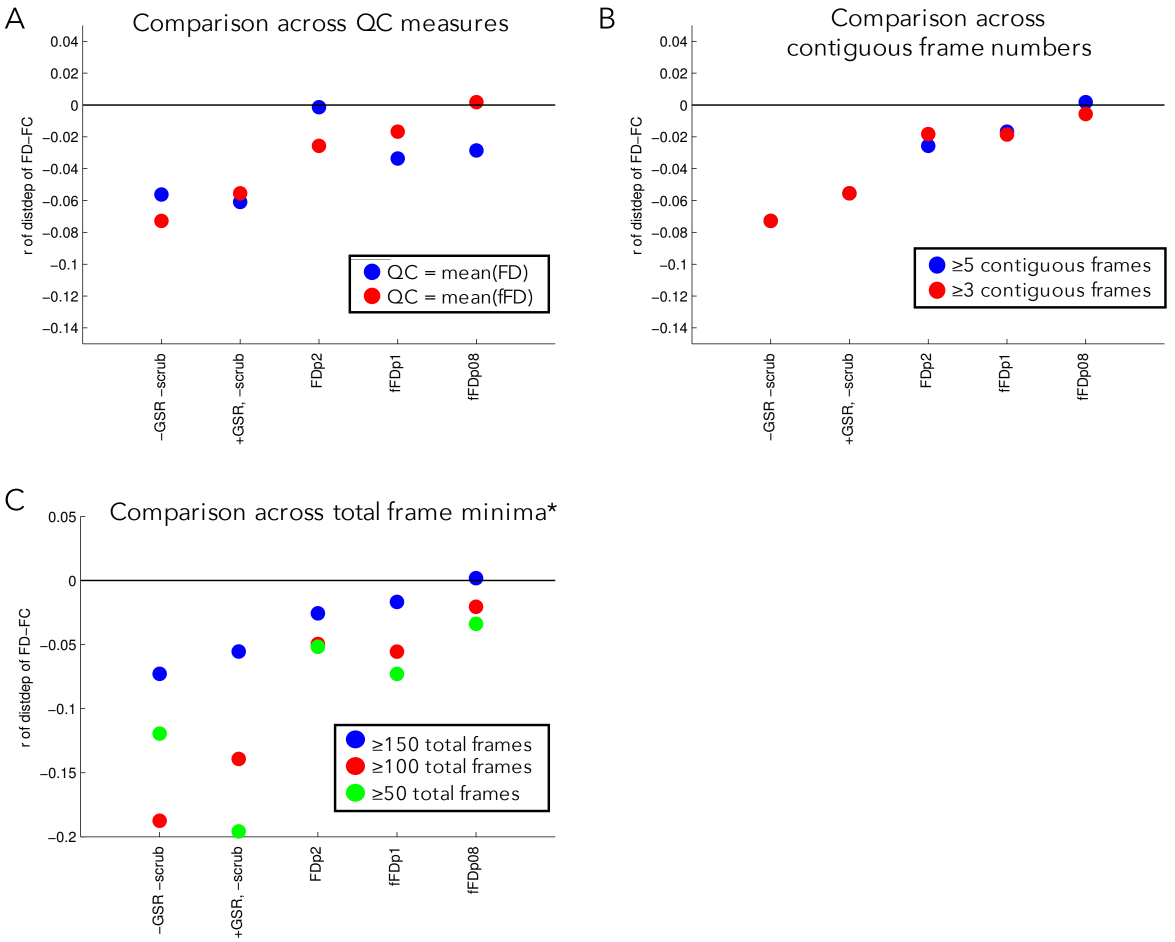


**Supplemental Figure 13:** QC-FC distance-dependence with other variations of analysis. (A) Comparison of the QC-FC distance dependent correlation (y-axis) under different preprocessing strategies when either mean FD or mean fFD are used as the QC quality metric. (B) Comparison of the distance dependence when either 3 or 5 contiguous frames are required in a data segment. (C) Comparison of the distance dependence when different numbers of total frame minima are imposed on the data. This comparison shows large magnitude differences. *Note that slightly different subjects were included in the groups in C (150: N = 79 (52 PD, 27 HC); 100: N = 87 (60 PD, 27 HC); 50: N = 103 (73 PD, 30 HC)).

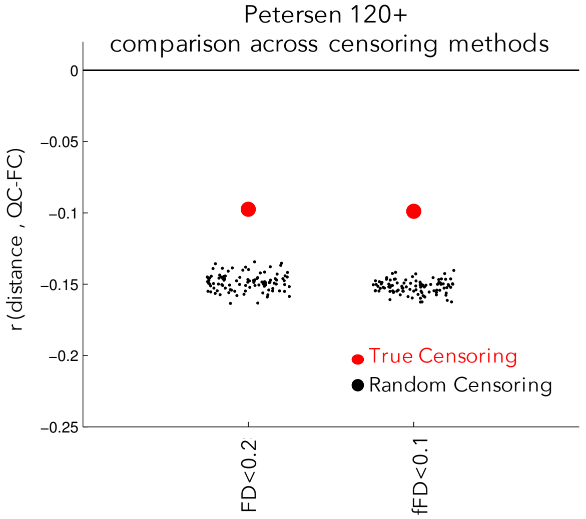


**Supplemental Figure 14: Distance dependence of QC-FC for different FD censoring approaches in the Petersen 120+ dataset.** The figure above shows the comparison between censoring with the FD<0.2 criteria vs. censoring with the fFD<0.1 criteria in N = 81 subjects from the Petersen-120+ dataset. As with PIB-PD dataset results reported in Figure 6, both censoring criteria do similarly at reducing distance dependence in QC-FC and outperform random censoring.

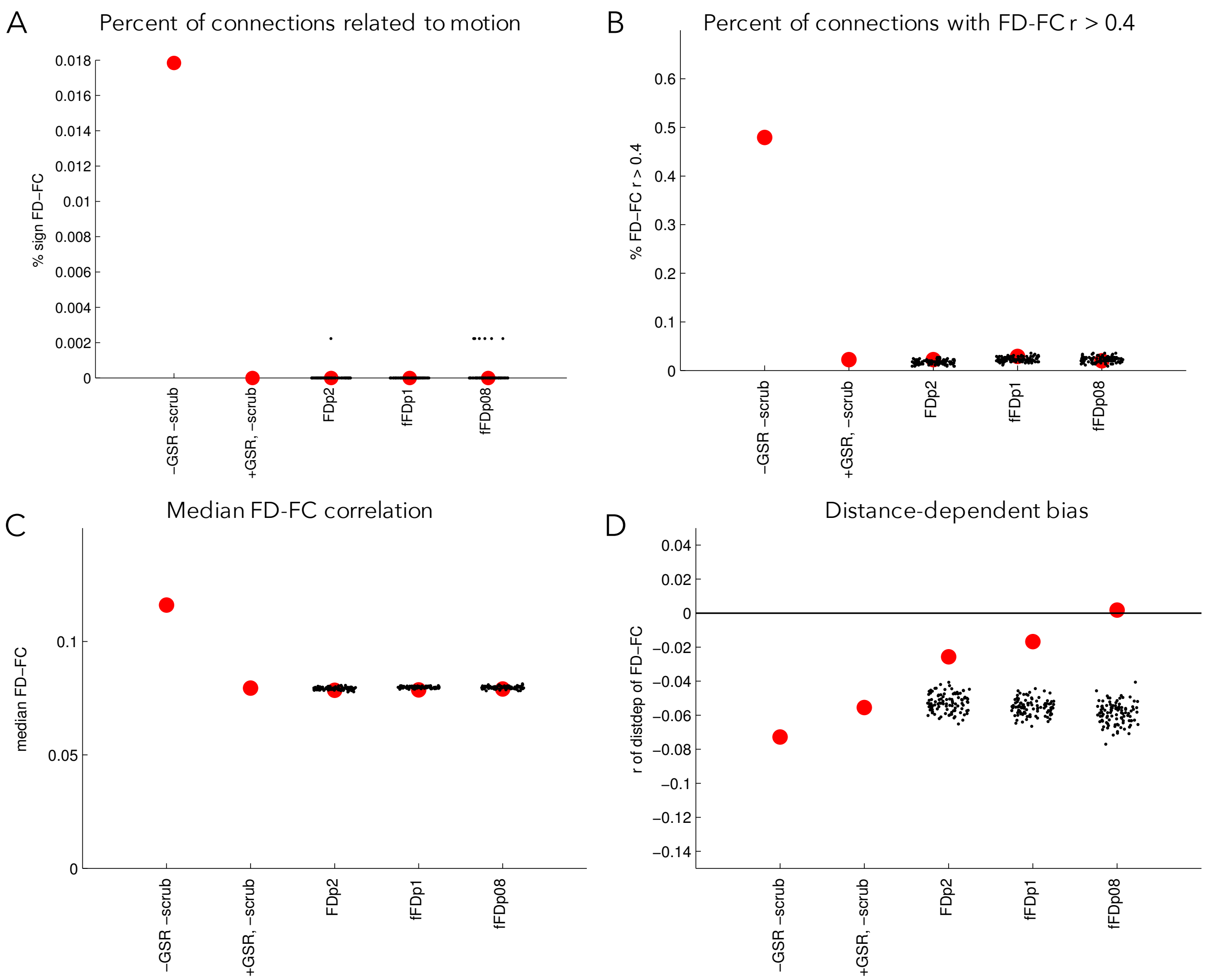


**Supplemental Figure 15:** Consequences of different censoring strategies on (A) the percentage of connections with significant relationships to motion, (B) the percentage of connections with a relationship to motion > 0.4, (C) the median FD-FC relationship, and (D) the distance-dependence of the FD-FC relationship, reproduced from Fig. 6 in the main paper. As can be seen, GSR primarily influences A-C, while censoring strategies have the strongest influence on D.

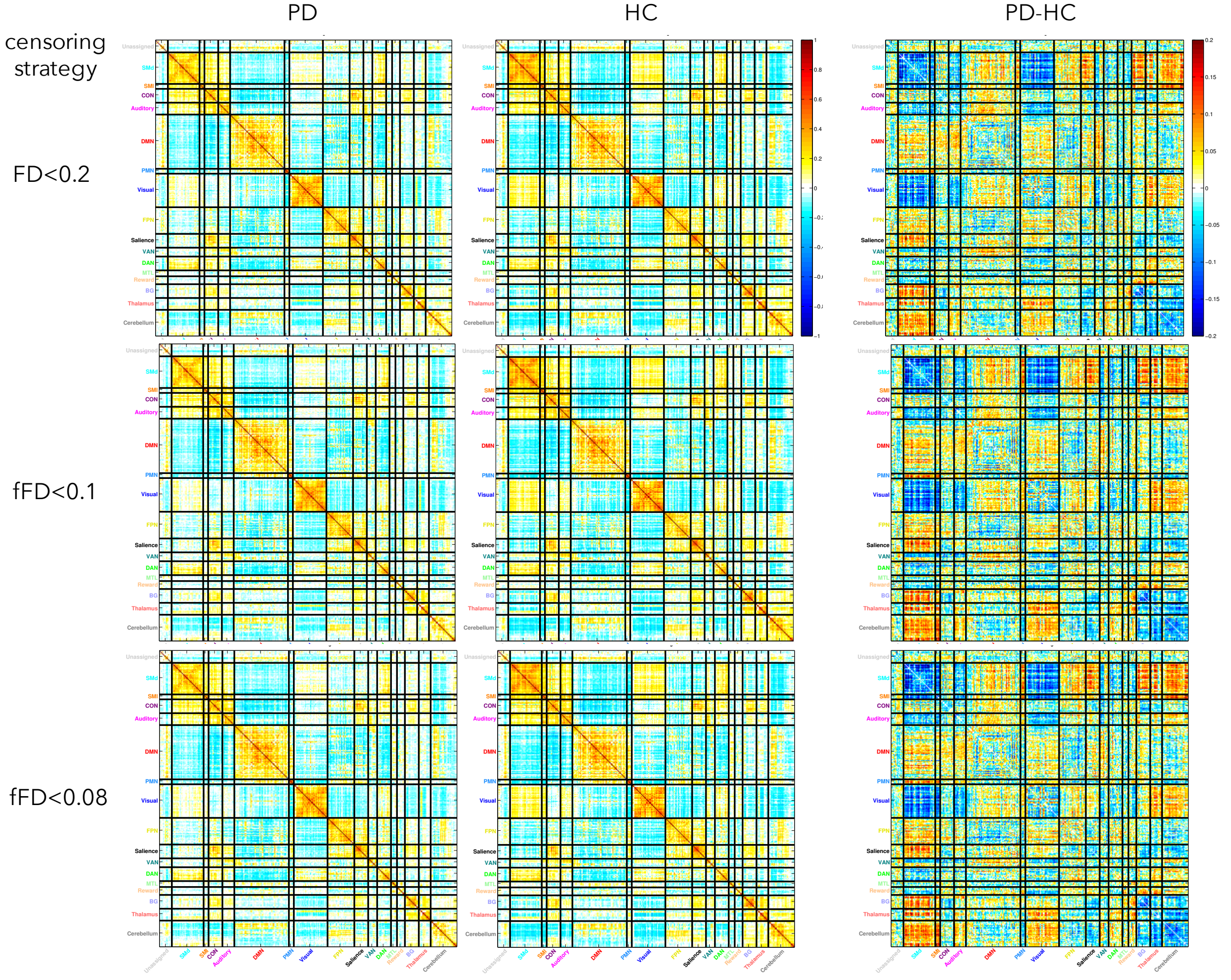


**Supplemental Figure 16:** Group average functional connectivity matrices for PD participants (left column), HC (middle column), and the difference between the two groups (right column) after processing with three different censoring strategies. These group functional connectivity results are quite similar regardless of the processing strategy employed. Note that the same participants were included in all three analyses, but the additional data savings from the fFD censoring strategies could lead to the inclusion of additional participants in analysis.
